# Vascular transcriptional and metabolic changes precede progressive intra-renal microvascular rarefaction in autosomal dominant polycystic kidney disease

**DOI:** 10.1101/2025.06.30.662386

**Authors:** Gizem Yilmaz, Santu K. Singha, Bansi Savaliya, Ahmed Abdelfattah, Walaa Elsekaily, Xiaohong Xu, Youwen Zhang, Christian Hanna, Marie C. Hogan, Alejandro R. Chade, Alfonso Eirin, Maria V. Irazabal

## Abstract

**Background:** The mechanisms contributing to progressive kidney damage in autosomal dominant polycystic kidney disease (ADPKD) remain unclear. Renal microvascular (MV) rarefaction plays an important role in kidney disease, but its natural history, underlying mechanisms, and contributions to renal disease progression in ADPKD remain unknown. We hypothesized that intrarenal MV rarefaction is present early on and is preceded by vascular transcriptional and metabolic changes.

**Methods:** *Pkd1*^RC/RC^ and WT mice (n=16 each) were studied at 1, 6, and 12 months. Total kidney volume (TKV) was measured *in vivo* (MRI), whereas renal MV architecture (3D-micro-CT), capillary density, perivascular fibrosis, and histomorphometric parameters were assessed *ex vivo*. In randomly selected *Pkd1*^RC/RC^ and WT kidneys (n=5, each/timepoint), mRNA-sequencing was performed to identify differentially expressed vasculature-related genes (DEGs). Next, in young individuals with ADPKD and matched controls (n=10 each), plasma cellular energy metabolites were determined (LC-MS/MS), validated in an extended cohort (n=32 and n=16, respectively), and correlated with markers of disease severity and progression. Gene–metabolite interaction networks were generated to integrate DEGs in *Pkd1*^RC/RC^ at 1 month with metabolites dysregulated in individuals with ADPKD.

**Results:** Renal MV density was preserved at 1 month but progressively decreased at 6 and 12 months, associated with capillary loss and perivascular fibrosis. A total of 110, 48, and 201 DEGs were identified at 1, 6, and 12 months, respectively. Plasma gamma-aminobutyric acid (GABA) and homocysteine (Hcy) levels were higher in individuals with ADPKD versus controls, interacted with DEGs implicated in inflammatory and innate immune response and Hcy metabolism, and correlated with TKV and renal blood flow.

**Conclusions:** Our data demonstrates that intrarenal MV abnormalities present early in ADPKD and are preceded by vascular transcriptional and metabolic changes. The renal microcirculation may constitute an important therapeutic target in ADPKD, and its underlying biomarkers may serve to monitor its progression.

**TRANSLATIONAL STATEMENT:** We provide the first longitudinal and most comprehensive analysis of the intrarenal microvascular network in a slowly progressive orthologous model of ADPKD and integrate the findings with studies in a young cohort of ADPKD individuals. Our integrated preclinical and clinical data identify vasculature-related pathways that could be targeted for therapeutic interventions and contribute promising, noninvasive biomarkers in patients with ADPKD. Furthermore, because alterations of the intrarenal microcirculation may affect drug delivery, a better understanding of its longitudinal changes may aid in treatment management.

## INTRODUCTION

Autosomal Dominant Polycystic Kidney Disease (ADPKD) is an important cause of kidney failure (KF) among adults, affecting up to 12.5 million people worldwide^1^. Currently, treatment options are limited to one FDA-approved drug (tolvaptan), which slows kidney function decline but does not stop disease progression^2,3^. In addition, tolvaptan-treated patients may experience aquaresis-related adverse events and hepatotoxicity in rare cases^3–6^. Understanding the mechanisms contributing to kidney damage is essential to identifying additional potential therapeutic targets and biomarkers for the disease.

The renal microcirculation is a highly organized system that transports oxygen and nutrients and removes toxins and waste products to maintain the kidney’s functional and structural integrity^7,8^. Importantly, microvascular (MV) dysfunction, damage, and loss - collectively referred to as MV rarefaction-play an important role in the progression of kidney disease^9^. Abnormal renal MV architecture, capillary loss, and angiogenesis have been reported in rodent models of PKD^10–13^. Likewise, in individuals with ADPKD, the vessels surrounding cysts are tortuous, abnormally patterned, and dilated^14,15^. However, these studies were performed in autosomal recessive models^10,11,13^ or autosomal dominant but with limited follow-up^12^, and in individuals in the late stages of the disease^14,15^. Thus, the natural history of renal MV rarefaction and its contributions to renal disease severity and progression in ADPKD remain unknown.

The plasticity of the renal microvasculature refers to its ability to modify the number, shape, and function of vessels to adapt to changes in the kidney microenvironment^8^. The vascular endothelium modulates many aspects of vascular function (VF), including control of vascular tone, permeability, proliferation, and fluid balance^16,17^. Endothelial dysfunction (ED), an imbalance between vasodilating (particularly nitric oxide, NO) and vasoconstricting substances, is an early event in MV disease^18^ and has been implicated in several diseases, including ADPKD^19,20^. In ADPKD, ED can precede hypertension (HTN) and KF, suggesting an underlying primary defect^19,20^. A key element associated with ED in ADPKD constitutes a reduction in NO, possibly due to defective ciliary-sensory mechanisms^21,22^, and/or decreased constitutive NO synthase (NOS)^19,20^. NOS may be also inhibited by elevated levels of asymmetric dimethylarginine (ADMA)^23–27^ resulting from reduced dimethylarginine dimethylaminohydrolases (DDAH) expression/activity through increased homocysteine (Hcy)^26,28–34^. NO levels may decrease also by oxidative degradation by Hcy^29,30^, or increased reactive oxygen species (ROS)^11,25,26^. Nevertheless, the early mechanisms associated with ED in ADPKD remain unknown.

Studies in individuals with ADPKD have assessed VF noninvasively in conduit vessels and using biochemical markers of ED^35–40^. Interestingly, in some of them^36,40^, changes in biochemical markers preceded differences in VF. These findings differed from studies in small-resistance vessels^19,20^, suggesting that ED in individuals with early ADPKD may be a particular feature of the microvasculature^25^. Contrarily, studies showed that individuals with ADPKD present with overt hemodynamic changes, evidenced by a decrease in renal blood flow (RBF), as an early vascular abnormality, parallelling total kidney volume (TKV) increase, preceding estimated glomerular filtration rate (eGFR) decline, and predicting structural and functional progression^41,42^. Overall, there is still a lack of knowledge on whether ED reflects micro/macrovascular damage and the sequence of events during the course of the disease. Importantly, an early, noninvasive biomarker reflective of underlying vascular pathology is still missing.

This study used a well-established slowly progressive mouse model of ADPKD (*Pkd1*^RC/RC^), state-of-the-art three-dimensional (3D) micro-computed tomography (micro-CT), high-throughput mRNA-sequencing (seq), and a young cohort of well-characterized, non-hypertensive, early-stage individuals with ADPKD, to comprehensively: **1)** characterize the longitudinal changes of the intrarenal MV from early stages, **2)** investigate molecular mechanisms associated with MV rarefaction, and **3)** identify early promising biomarkers in the plasma of individuals with ADPKD reflective of underlying vascular pathology. We hypothesized that intrarenal MV rarefaction is present from the early stages of the disease, and preceded by vascular transcriptional and metabolic changes, which can be detected early on in the plasma of individuals with ADPKD.

## METHODS

### Animal studies

*Pkd1*^RC/RC^ and WT mice (n=16/each, 8M/8F) were studied at 1, 6 and 12 months (IACUC#: A00003777-A00005742). All methods are reported in accordance with ARRIVE guidelines^43^.

Animals were scanned by magnetic resonance imaging (MRI) at each time-point, and TKV was estimated^44^ and adjusted by animals’ body length (blTKV). Blood pressure was monitored noninvasively. At euthanasia, kidney/body weights (KW/BW) were recorded, and plasma was collected for BUN. The left kidney was harvested^45^, and kept at -80°C for mRNA-seq or quantitative polymerase chain reaction (qPCR). The right kidney was prepared for histology.

Renal MV architecture was assessed in both kidneys using micro-CT in WT and *Pkd1*^RC/RC^ mice (3M/3F, 12 kidneys/time-point). After euthanasia, the kidneys were perfused with a radio-opaque silicone polymer (Microfil) until all vessels were filled^46–48^. The polymer-filled kidneys were allowed to cure and placed in formalin before scanning^49^. The number of renal microvessels of diameters 0-500μm in each section was calculated using Analyze and expressed as spatial density (number of vessels/tissue area, adjusted for cysts from MRI). The cortex and medulla were divided into outer-inner regions^50^ (OC, IC, OM, and IM), and MV density and diameter were calculated at each level. Tortuosity, an index of neovascularization and MV immaturity^51^ was calculated in tomographically isolated microvessels, as described^52^. Cystic (CA) and fibrotic areas (FA) were determined from kidney sections (n=2 or 8/animal) stained with hematoxylin-eosin and Picosirus-Red, respectively^11^. Peritubular capillary density was determined in H&E-stained sections (n=10/animal)^53^, further adjusted by CA, and expressed as the number of capillaries/parenchyma^11^. Perivascular fibrosis was determined in Masson’s trichrome-stained slides and expressed as % fibrosis/parenchyma.

In randomly selected WT and *Pkd1*^RC/RC^ mice (n=5/each/timepoint), mRNA-seq was performed at the Mayo Clinic GAC, as described^54,55^. The total RNA was extracted, RNA libraries were prepared, samples were sequenced, and processed^56^, and gene expression was quantified^57^ and normalized by Counts per Million (CPM). To visualize the whole spectrum of changes in the expression of vasculature-related genes in WT and *Pkd1^RC/RC^*kidneys at 1, 6, and 12 months, we used the Mouse Genome Informatics database to screen vasculature-related genes (**Table_S1**). Differentially expressed genes (DEGs) (CPM>0.1, fold-change>1.4 or <0.7, and p<0.05) were identified using edgeR^58^, FDR-corrected^59^, (**Supp_Data_files_1-3**) and classified based on their cellular component, molecular function, and biological process^60^, and interrogation of protein functional and physical interactions was performed^61^. Validation of mRNA-seq was performed in randomly selected top-upregulated (FC>2.5, p<0.001) and top-downregulated (FC<0.4, p<0.001) mRNAs in *Pkd1*^RC/RC^ versus WT kidneys at 1, 6 and 12 months by qPCR^62^, normalized to β-actin (*Actb*).

### Human studies

The study was approved by Institutional Review Board of the Mayo Clinic (IRB 24-013591) in accordance with the Declaration of Helsinki and the Health Insurance Portability and Accountability Act guidelines. Thirty-two, young individuals (8-35 years of age), with SBP<130mmHg without antihypertensive medication (individuals>13 years of age) or <95th%ile for height, age and gender (individuals<13 years of age)^63^, with eGFR>90mL/min/1.73m², and a previous diagnosis of ADPKD (Ravine/Pei criteria)^64,65^ were matched 2:1 for age (±2 years)/gender to control individuals without a personal or family history of kidney disease and included in this cross-sectional study. Exclusion criteria included atypical presentation^66^, antihypertensive medication, diabetes mellitus, predicted urine protein excretion>1g/24hrs, abnormal urinalysis, a systemic disease known to affect the kidney, or a known pathogenic variant in other than in *PKD1* or *PKD2* genes.

Blood and urine samples were collected, and an abdominal MRI/wo contrast was performed to estimate TKV (n=28/ADPKD, n=14/controls) and RBF (n=21/ADPKD, n=14/controls), as described^67–69^. TKVs were adjusted by the patient’s height (htTKV) and RBFs by BSA. Patients in which htTKV was available (n=28), were classified based on their htTKV/age^66^. The eGFR was calculated with the CKD in Children (CKiD) equation for individuals<18 years old and with the CKD-epi equation (2021) for individuals>18 years^70,71^. Genetic data was available in 19 individuals with ADPKD.

Plasma concentrations of metabolites involved in cellular energy metabolism including tricarboxylic acid cycle (TCA), glycolytic pathway, fatty acids (FAs) and amino acids (AAs) were determined by liquid chromatography-tandem/mass spectrometry (n=10/ADPKD, n=10/matched-controls)^72–78^.

Gene–metabolite interaction networks were generated using MetaboAnalyst by mapping vasculature-related DEGs in *Pkd1*^RC/RC^ at 1 month and metabolites dysregulated in individuals with ADPKD^79^.

Plasma Hcy, gamma-aminobutyric acid (GABA) and ADMA levels were measured by enzyme-linked immunosorbent assay (ELISA) in the full cohort (n=32/ADPKD, n=16/matched-controls).

### Statistical analysis

Statistical analysis was performed using PRISM9. The Shapiro-Wilk test was used to test for deviation from normality. Parametric (Student’s t-test) and nonparametric (Kruskal-Wallis) tests were performed when appropriate. p-values ≤0.05 were considered significant.

Extended methods are available as supplemental material.

## RESULTS

### Animal Characteristics

Animal characteristics at each time-point are shown in **Table_1 and Figure_S1**. BW and urine output were similar between WT and *Pkd1*^RC/RC^ mice. However, *Pkd1*^RC/RC^ mice exhibited higher KW/BW, blTKV **(Figure_S1A)**, and CA **(Figure_S1B)** at all time-points. On the other hand, FA **(Figure_S1C)** and BUN were not different at 1 and 6 months but were higher in *Pkd1*^RC/RC^ at 12 months. SBP, DBP, and MAP pressure did not differ at 1 month but were higher in *Pkd1*^RC/RC^ at 6 and 12 months. In *Pkd1*^RC/RC^ mice, KW/BW, blTKV, and CA increased from 1 to 6 months, in agreement with previous observations in this model^80^. Yet, only blTKV and CA further increased at 12 months. FA and BUN did not differ between 1 and 6 months, but increased from 6 to 12 months, consistent with the progression of the disease observed in patients with ADPKD^81–83^.

### MV spatial density and diameter are preserved early on, but change as the disease progresses, starting in the cortical region

The average spatial density of intra-renal microvessels was similar between *Pkd1*^RC/RC^ and WT mice at 1 month, markedly decreased in *Pkd1*^RC/RC^ at 6 months, and decreased further at 12 months (**Figure_1A**). Similarly, the average vessel diameter was not different at 1 month in *Pkd1*^RC/RC^ versus WT but was higher at 6 and 12 months.

**Figure 1.**
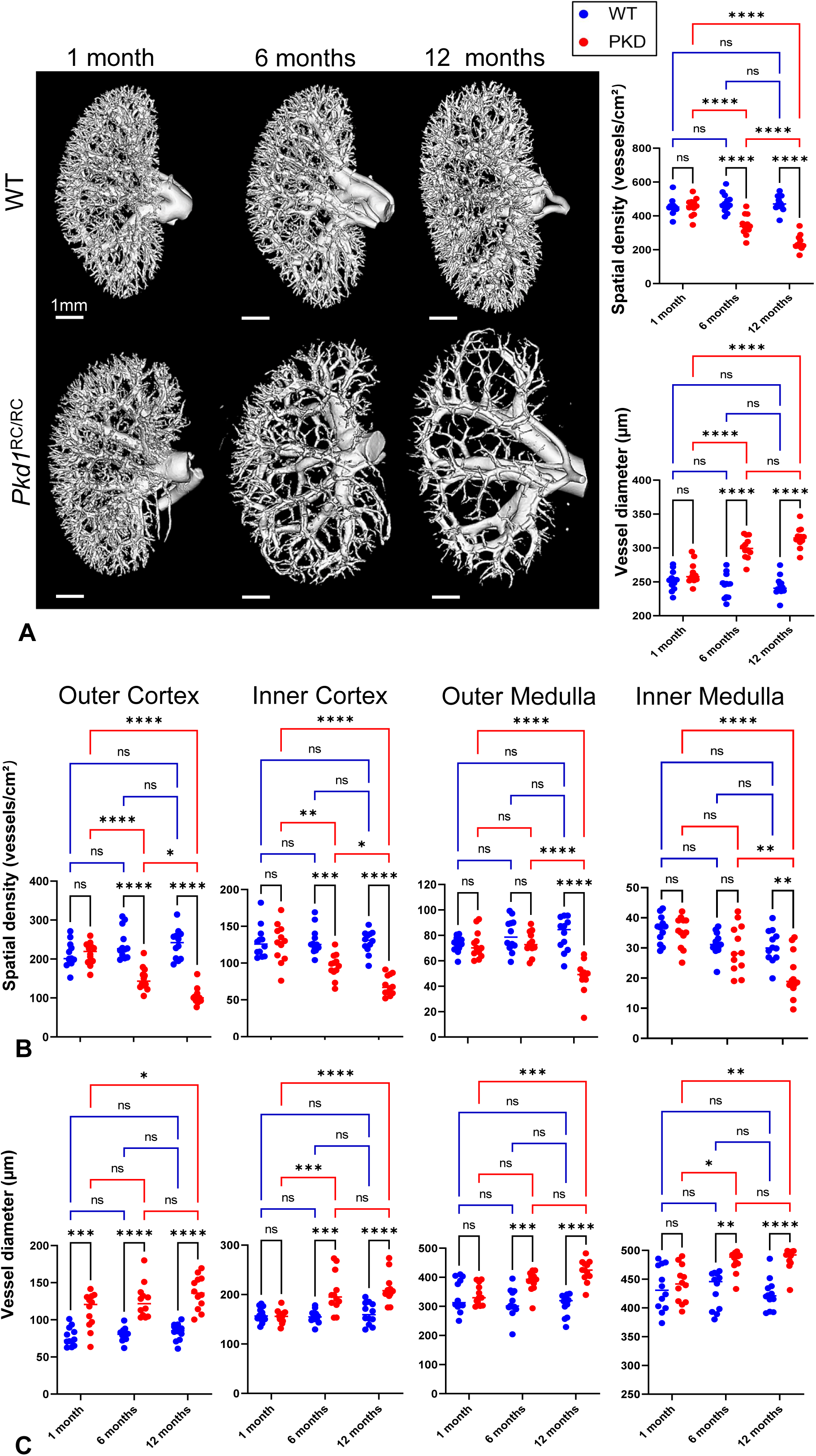
Renal microvascular rarefaction in Pkd1^RC/RC^. Representative 3-dimensional micro-computed tomography (micro-CT) images of the kidney of *Pkd1^RC/RC^*and WT mice at 1, 6, and 12 months, and quantification of microvascular spatial density and average vessel diameter in the whole kidney **(A)**. Quantification of microvascular spatial density **(B)** and average vessel diameter **(C)** in the outer cortex, inner cortex, outer medulla, and inner medulla (n=12/genotype). *p<0.05, **p<0.01, ***p<0.001, and ****p<0.0001 compared to WT at each timepoint.

When analyzed by regions (OC, IC, OM, and IM), MV spatial density followed a similar pattern as the average and was not different between *Pkd1*^RC/RC^ and WT mice at 1 month (**Figure_1B**). At 6 months, the OC and IC regions exhibited a decrease in MV density, whereas the OM and IM maintained a similar density in *Pkd1*^RC/RC^ compared to WT mice. By 12 months, all regions of *Pkd1*^RC/RC^ kidneys presented with a decrease in MV density compared to WT.

The OC vessel diameter was higher in *Pkd1*^RC/RC^ versus WT at 1 month and remained similarly elevated at 6 and 12 months, whereas the IC vessel diameter remained at WT levels at 1 month and increased and remained higher at 6 and 12 months (**Figure_1C**). Both outer and inner medullary regions presented comparable vessel diameters in *Pkd1*^RC/RC^ and WT at 1 month, but vessel diameters were higher in *Pkd1*^RC/RC^ versus WT in both regions by 6 and 12 months.

### Increased MV tortuosity is an early sign of MV structural abnormalities in Pkd1^RC/RC^

Cortical peritubular capillary density did not differ between *Pkd1*^RC/RC^ and WT at 1 month, decreased in *Pkd1*^RC/RC^ at 6 months, and further decreased at 12 months (**Figure_2A**), in line with the changes in the renal MV density.

**Figure 2.**
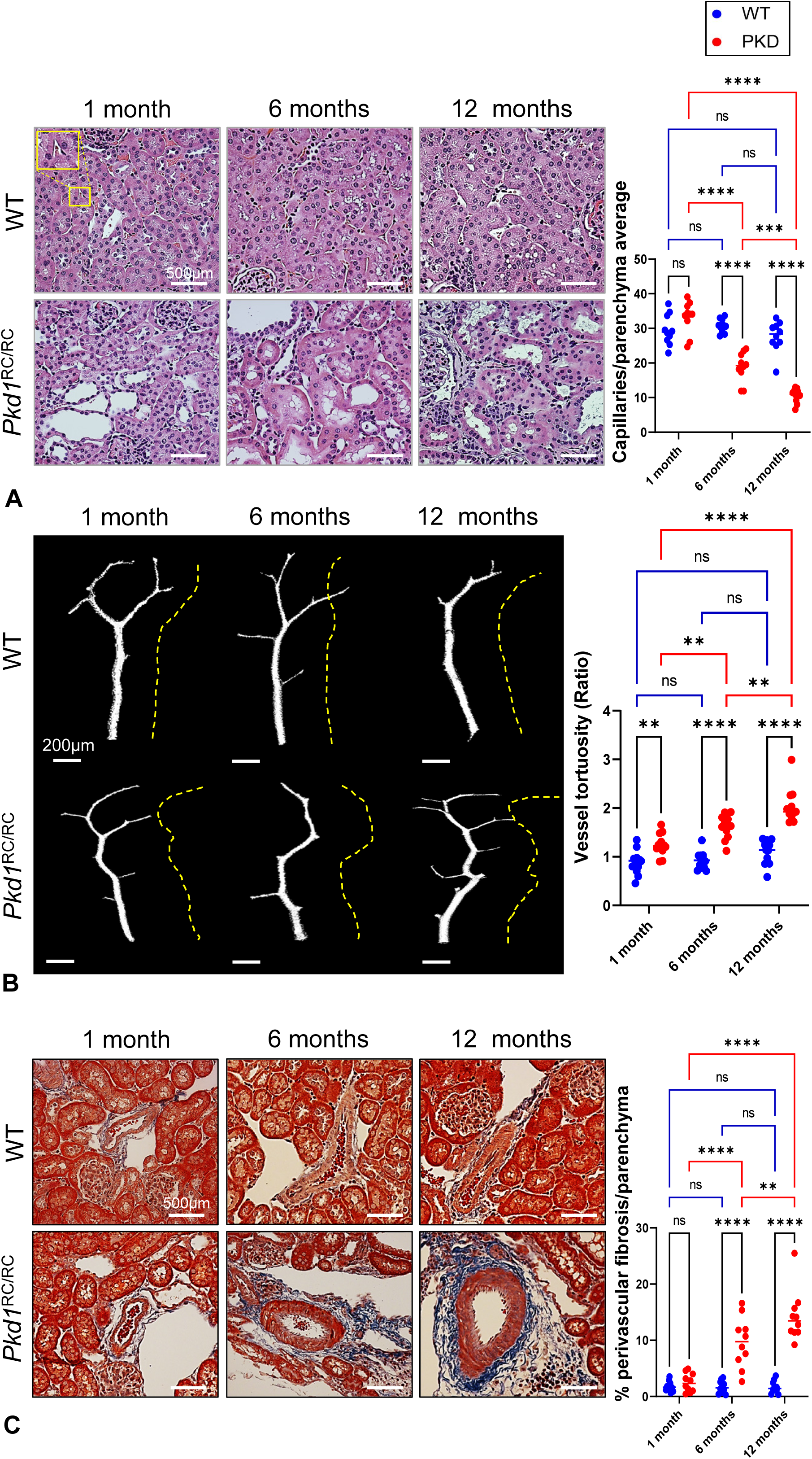
Renal capillary loss, microvascular remodeling and perivascular fibrosis in Pkd1^RC/RC^. Representative hematoxylin and eosin (H&E)-stained kidneys (40X images) and quantification of the number of capillaries per field/parenchyma area in *Pkd1^RC/RC^* and WT kidneys at 1, 6, and 12 months (n=5M and 5F per genotype for each timepoint) **(A)**. Tomographically isolated vessels (3D-micro-CT) and quantification of their tortuosity (n=12/genotype). Dashed yellow lines indicate the entire vascular path length (actual length) of each microvessel (from base to tip) **(B)**. Masson’s trichrome staining of *Pkd1^RC/RC^* and WT mouse kidneys at 1, 6, and 12 months and quantification of perivascular fibrosis and expressed as % fibrosis/parenchyma (n=5M and 5F per genotype for each timepoint) **(C)**. **p<0.01, ***p<0.001, and ****p<0.0001 compared to WT at each timepoint.

Because capillary regression may be masked by concomitant angiogenesis, creating an overall appearance of a normal capillary density, we determined MV tortuosity, an index of neovascularization, and MV immaturity^51^. Vessel tortuosity was higher in *Pkd1*^RC/RC^ versus WT at 1 month, increased at 6 months, and increased further at 12 months, suggesting progressive vascular wall remodeling from the early stages of the disease (**Figure_2B**).

Perivascular fibrosis, characterized by increased connective tissue deposition (particularly collagen), around the vessels, has been demonstrated in the kidney and is a hallmark of vascular pathology^84,85^. At 1 month, perivascular fibrosis was not different in *Pkd1*^RC/RC^ versus WT. Interestingly, although global kidney fibrosis was not different from WT until 12 months **(Figure_S1C)**, perivascular fibrosis was higher in *Pkd1*^RC/RC^ at 6 months and progressively increased at 12 months (**Figure_2C**).

### mRNA-seq and functional clustering analysis identify molecular features and biological processes accompanying the vascular changes at different stages of the disease

To identify the molecular features accompanying the observed structural changes, we performed vasculature-related gene (**Table_S1**) differential expression analysis at the same time-points. We identified 101, 39 and 167 genes upregulated and 9, 9 and 34 genes downregulated in *Pkd1*^RC/RC^ compared to WT at 1, 6 and 12 months, respectively (**Figure_3**).

**Figure 3.**
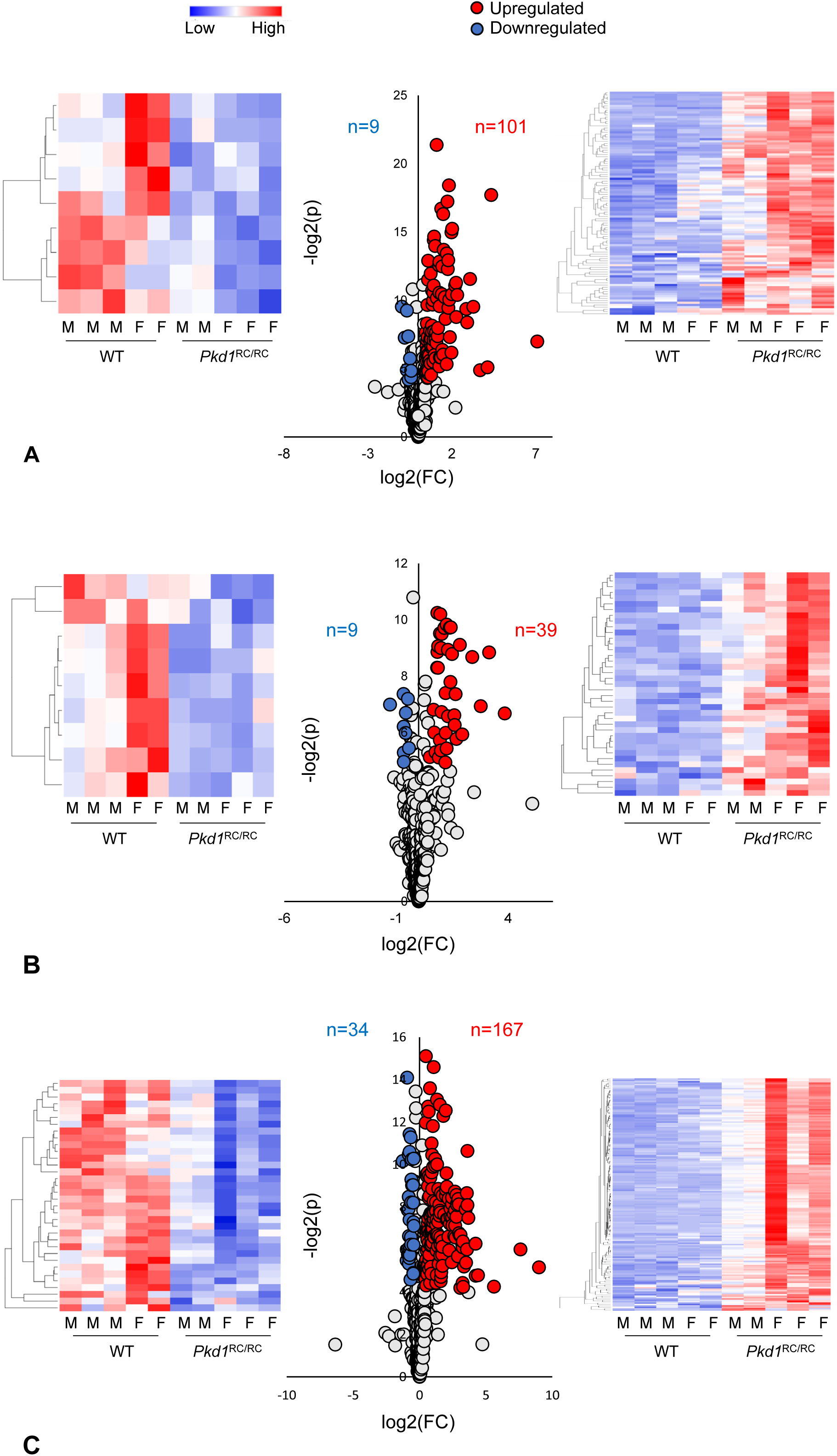
Longitudinal changes in the expression of vasculature-related genes in Pkd1^RC/RC^ kidneys. Volcano plot and heat maps showing vasculature-related genes upregulated and downregulated in *Pkd1*^RC/RC^ compared to WT kidneys at 1 **(A)**, 6 **(B)**, and 12 months **(C)**. The vertical axis (y-axis) corresponds to the -log 2 (p-value) and the horizontal axis (x-axis) displays the log 2-fold change (*Pkd1*^RC/RC^/WT) value. Upregulated and downregulated genes are shown as red and blue dots, respectively, while non-significant genes are shown as gray dots. (n=3M and 2F WT and n=2M and n=3F *Pkd1^RC/RC^* for each timepoint).

To validate our mRNA-seq analysis, we performed qPCR on randomly selected candidate genes at each time-point and found that their expression followed the same patterns as the mRNA-seq **(Figure_S2).**

To gain further insights into the biological processes associated with vascular structural changes, we performed functional clustering analysis of DEGs. Vasculature-related genes upregulated in *Pkd1*^RC/RC^ at 1 month encode proteins primarily implicated in vasculature development, blood vessel morphogenesis, cell motility, and response to endogenous stimulus (**Figure_S3**). Genes downregulated in *Pkd1*^RC/RC^ at 1 month produce proteins involved in vascular development and blood vessel morphogenesis, and cell specialization and angiogenesis involved in wound healing (**Figure_S4**).

Genes upregulated in *Pkd1*^RC/RC^ at 6 months generate proteins implicated in vasculature development and blood vessel morphogenesis, cell adhesion, motility and chemotaxis (**Figure_S5**). However, genes downregulated in *Pkd1*^RC/RC^ at 6 months encode proteins involved in endothelial cell proliferation, tyrosine kinase signaling and sprouting angiogenesis (**Figure_S6**).

Genes upregulated in *Pkd1*^RC/RC^ at 12 months generate proteins involved in vasculature-related morphogenesis, cell adhesion and migration (**Figure_S7**). Genes downregulated in *Pkd1*^RC/RC^ at 12 months mainly encode proteins involved in blood vessel morphogenesis, vasculature development and endothelial cell migration (**Figure_S8**).

### Pilot metabolomics analysis from individuals with ADPKD and matched controls identifies GABA and Hcy as metabolites of interest early on the disease

The lack of human tissue samples from children and young adults, in which renal biopsies are not routinely indicated and nephrectomies are not performed until KF, prevents the identification of vascular changes early on noninvasively. The mammalian kidney is continuously active with marked metabolic activity and high energy demands^86^. Importantly, studies show that vascular structural changes in the kidney may result from an adaptation and/or response to changes in metabolic demands^8^. Metabolomics analyses can be utilized to determine multiple metabolite concentrations in a particular tissue or biofluid from a living organism^87^. Therefore, to noninvasively assess early metabolic changes in individuals with ADPKD that may be associated with vascular abnormalities, we performed a pilot study to determine plasma levels of metabolites involved in cellular energy metabolism, including the TCA cycle, glycolytic pathway, FAs and AAs.

**Table_S2** summarizes the clinical, laboratory, and demographic characteristics of the individuals included in the pilot study. Individuals with ADPKD and controls had an average age of 23.1±3.1 and 22.7±3.1 years, respectively. None of the biometrics, clinical or laboratory parameters differed between the groups. As expected, htTKV was higher in patients with ADPKD. Three patients were classified as class 1B, four as 1C, and three as 1D^66^. Analysis of plasma samples identified and quantified 54 metabolites **(Table_S3)**, of which only 2, GABA and Hcy, were different between the groups (pl*<*l0.05). Therefore, plasma GABA and Hcy emerged as metabolites of interest early on the disease.

### Multi-omics integration analysis reveals two key vasculature-related networks at the early stages of the disease

Plasma metabolites may primarily reflect changes in the systemic compartment and may not be related to the molecular features accompanying the observed intrarenal MV structural changes. Therefore, to visualize GABA and Hcy within the context of biological processes that may be associated with the observed intrarenal MV changes, we performed an integrated analysis using the DEGs identified in the tissue from *Pkd1*^RC/RC^ animals at 1 month.

The integration of GABA and Hcy into the DEGs resulted in 2 deregulated networks at both the transcript and metabolite level, and containing at least 3 nodes (**Figure_4**). Specifically, this interaction links GABA to inflammation and components of the innate immune system genes *Cxcl10*, *Anxa1*, *C3*, *C3ar1*, *C5ar1*, *Dcn,* and *Ackr3*, all of which were upregulated in *Pkd1*^RC/RC^ animals (**Figure_4A**). Hcy was not only linked to *Ddah1* and *Gpx1*, which are directly related to its metabolism and were downregulated in *Pkd1*^RC/RC^ animals, but also to *Serpine*, *Mmp2*, *Ccl2*, and *Ccr2*, which were upregulated in *Pkd1*^RC/RC^ animals and are implicated in tissue injury repair, remodeling and angiogenesis as well as inflammation and monocyte chemotaxis (**Figure_4B**).

**Figure 4.**
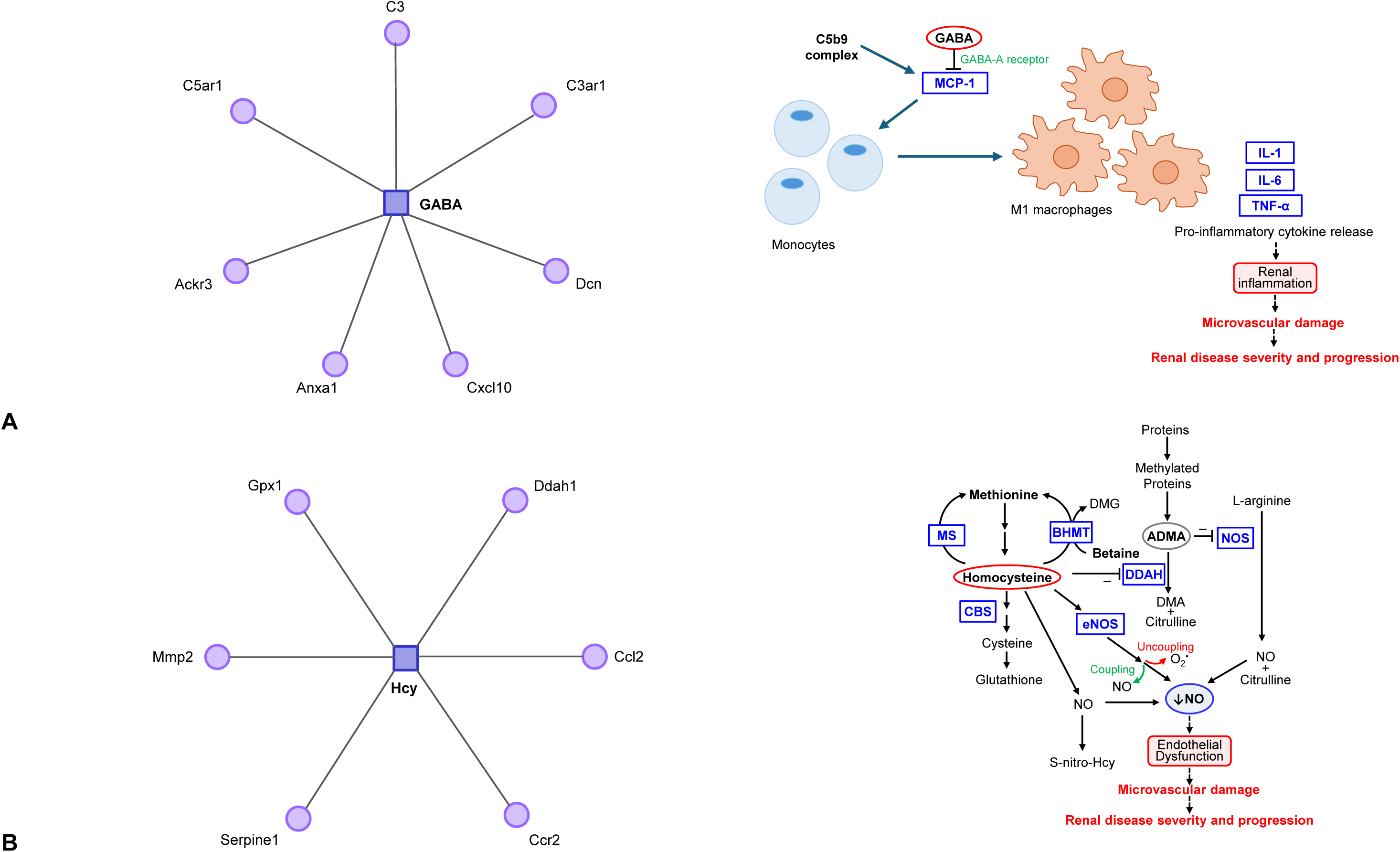
Multi-omics integration analysis reveals two key vasculature-related networks at the early stages of the disease.in ADPKD. Gene–metabolite interaction networks (MetaboAnalyst) of differentially expressed vasculature-related genes in *Pkd1*^RC/RC^ at 1 month (circles) and circulating levels of gamma-aminobutyric acid (GABA) **(A)** and homocysteine (Hcy) **(B)** (squares) in young individuals with ADPKD (left). Schematic diagram showing the potential role of GABA and inflammation and the role of endothelial dysfunction and microvascular damage (right).

### GABA, Hcy and ADMA are promising vasculature-related biomarkers from the early stages of the disease

To establish the role of GABA and Hcy as vasculature-related biomarkers in the early stages of the disease, we extended our studies to a larger cohort of 32 individuals with ADPKD and 16 matched controls. In addition, because studies have shown that increased levels of Hcy lead to DDAH decreased expression/activity resulting in elevated ADMA^26,29–34^, and our findings in animals of reduced *Ddah1* expression, we determined plasma ADMA levels in the same groups. **Table_2** summarizes the clinical, laboratory, and demographic characteristics of the entire cohort. Individuals with ADPKD and controls had an average age of 21.4±5.6 and 22.3±5.4 years, respectively. None of the biometrics, clinical or laboratory parameters differed between the groups. As expected, htTKV was higher in patients with ADPKD. However, RBF was similar between the groups. Eleven patients were classified as class 1B, eleven as 1C, four as 1D, and two as class 1E^66^. The genotype was *PKD1* in 16 patients (84.2% of the 19 patients with genetic data available) and *PKD2* in 2 (10.5 %), whereas no mutation was detected in 1 patient (5.3 %). The plasma levels of GABA, Hcy and ADMA were higher in individuals with ADPKD compared to controls **(Figure_5A-C).**

**Figure 5.**
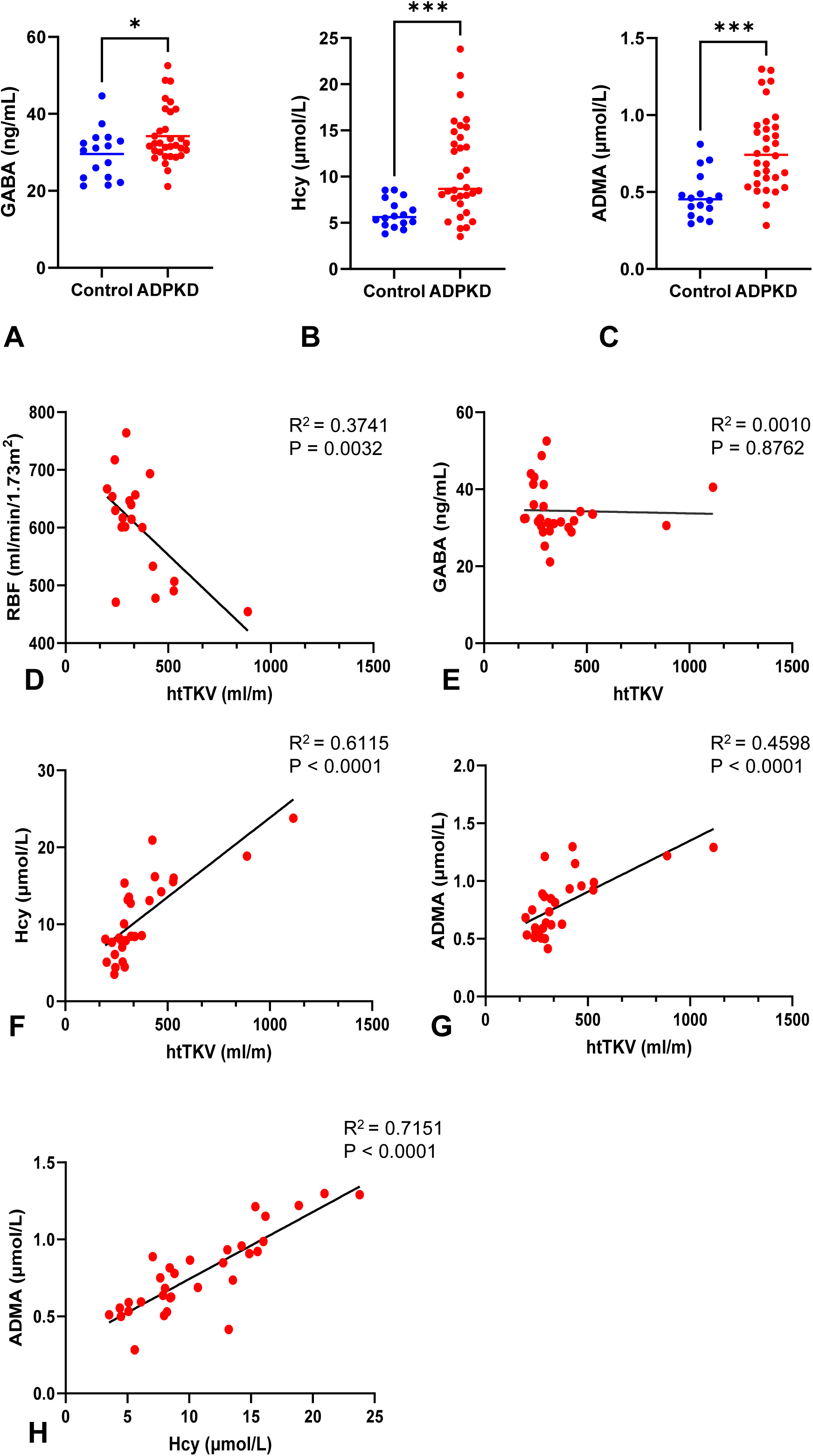
Plasma gamma-aminobutyric acid (GABA), homocysteine (Hcy), and asymmetric dimethylarginine (ADMA) levels in individuals with ADPKD and their correlation with markers o disease severity. Plasma GABA **(A)**, Hcy **(B)**, and ADMA **(C)** levels (ELISA) in individuals with ADPKD and age and sex-matched controls (n=32 and n=16, respectively). Correlations between renal blood flow (RBF) and height-adjusted total kidney volume (htTKV) **(D)**, GABA and htTKV **(E)**, Hcy and htTKV **(F)**, ADMA and htTKV **(G)**, and ADMA and Hcy **(H)**.

In individuals with ADPKD, although RBF was not different compared to controls, there was an inverse correlation with htTKV **(Figure_5D).** Contrarily, plasma GABA did not correlate with htTKV **(Figure_5E)**, RBF, or eGFR **(Figure_S9_A-B)**. Plasma Hcy strongly correlated directly with htTKV **(Figure_5F)** and had a weak inverse correlation with RBF (**Figure_S9C)** but no correlation with eGFR (**Figure_S9D**). Similarly, there was a direct correlation between plasma ADMA and htTKV **(Figure_5G)** and a weak inverse correlation with RBF **(Figure_S9E)** but no correlation with eGFR **(Figure_S9F).** Finally, there was a strong direct correlation between plasma Hcy and ADMA in individuals with ADPKD **(Figure_5H)**.

## DISCUSSION

We provide the first longitudinal and most comprehensive analysis of the intrarenal MV network in a slowly progressive orthologous model of ADPKD and integrate the findings with studies in a young cohort of ADPKD individuals. Our integrated preclinical and clinical data identify vasculature-related pathways that could be targeted for therapeutic interventions and contribute promising, noninvasive biomarkers in patients with ADPKD. Furthermore, because alterations of the intrarenal microcirculation may affect drug delivery, a better understanding of its longitudinal changes may aid in treatment management.

We found preservation of MV and peritubular capillary density at the early stages of the disease. As the disease progressed, capillary and MV loss and increased vessel diameter became apparent, starting from the cortical regions. Further analysis identified increased intrarenal MV tortuosity, suggestive of abnormal angiogenesis^51,88^, as the earliest sign of structural abnormalities. Perivascular fibrosis was higher in *Pkd1*^RC/RC^ at an earlier age than global kidney fibrosis, suggesting that early alterations in intrarenal microvessels might play a role in initiating renal fibrosis. This is in line with previous studies in kidney fibrosis models, demonstrating the central role of peritubular endothelial cells in the development of renal fibrosis^89^.

Our study identified molecular changes accompanying the architectural remodeling throughout the disease. At the early stages, we observed predominantly upregulation of genes implicated in vascular development and blood vessel morphogenesis, which may explain the preservation of the capillary density and vessel immaturity. These changes were present in the absence of a prominent cystic phenotype, suggesting that intrarenal vascular changes are not just a consequence of cystic displacement or disease progression. With disease progression, the downregulation of genes involved in vasculature development and blood vessel morphogenesis became more pronounced, in line with our structural analysis, showing a progressive decrease in MV and capillary density. Standing upregulated vasculature-related processes with disease progression were cell adhesion, inflammatory chemotaxis, and cell migration, consistent with the phenotypic changes observed in this model and in humans^80,90^. Other processes downregulated early on are related to angiogenesis in wound healing, whereas endothelial cell migration, which appears later on, may underlie abnormal repair mechanisms and contribute to its progression.

Various metabolic changes that can affect the intrarenal microvasculature have been described in PKD^91–97^. However, most reported changes are from tissue analyses. In humans, the lack of tissue samples at early-stages hinders the identification of early biomarkers underlying disease mechanisms. Plasma metabolomics offers a noninvasive systemic snapshot of the body’s overall metabolic state. Despite the very early-stage of our pilot population, by controlling for the feeding state and circadian cycle, known biological factors contributing to the highest variability^98^, performed under rigorous conditions, and gender/age-matched, we identified GABA and Hcy as biomarkers of interest. Further, our multi-omics approach provided a multi-layered insight into how upstream changes in the kidney may exert changes on downstream metabolic pathways affecting the intrarenal microvasculature. Specifically, we found that increases in the inflammatory and innate immune response, possibly linked through GABA, and abnormalities in Hcy metabolism may be important molecular features accompanying the intrarenal MV structural changes at the early stages.

Studies showed that GABA exerts protective effects in chronic kidney injury^99–101^, by modulating monocyte chemoattractant protein-1. Complement-related release of this cytokine triggers monocytes and pro-inflammatory macrophage activation, contributing to renal inflammation and MV damage^102^. Therefore, early elevations in plasma levels of GABA may reflect a compensatory mechanism to mitigate the inflammatory response in the PKD kidney. Alternatively, increased circulating GABA may reflect a dysregulation in the renal GABAergic system^103^, possibly due to abnormalities in its metabolism or to compensate for enhanced renal sympathetic nerve activity observed in patients with ADPKD^104–106^. Future mechanistic studies are needed to determine the role of GABA in ADPKD.

We and others have shown that in PKD, ED develops early on, prior to the development of HTN and KF^11,19,20^. However, the mechanisms are still unclear and likely multifactorial. Studies showed elevated levels of Hcy, ADMA and oxidative stress in hypertensive individuals with ADPKD and eGFR>60 ml/min/1.73/m^225,26^. The current study extends these observations in young individuals with ADPKD and preserved kidney function, demonstrating that circulating Hcy and ADMA are elevated before the development of HTN and overt hemodynamic changes. Elevated levels of Hcy have been associated with impaired endothelial-dependent NO-mediated vasodilatation and are an accepted marker of ED^31,107–110^. Hcy may decrease NO availability by oxidative degradation^29,30^, eNOS uncoupling, and NOS inhibition by increased ADMA, resulting from reduced DDAH expression/activity^29,32^. This study revealed a network interaction between elevated levels of Hcy in young individuals with ADPKD, decreased expression of *Ddah1* in kidneys from early-stage *Pkd1*^RC/RC^ animals and higher levels of ADMA in patients, suggesting a mechanism underlying vascular dysfunction at the early stages. Importantly, these changes occur prior to marked vascular structural changes, and position Hcy and ADMA as promising, noninvasive vascular-related early biomarkers in ADPKD.

A reduction in RBF is an early functional defect in ADPKD that parallels TKV increase, precedes eGFR decline and can be observed before the development of HTN^68,69^. In this early-stage cohort, RBF was not different from controls. This may be partially explained by the young population, the low number of patients, or a latency between molecular changes underlying ED and a reduction in RBF. Furthermore, it is in line with prior studies showing alterations in biochemical markers of ED preceded functional changes in conduit vessels^36,40^. Together with the intrarenal structural and molecular changes observed in our early-stage animals, and prior studies in small resistance vessels from individuals with ADPKD^19,20^, further suggests that in ADPKD, ED precedes overt hemodynamic changes and compromises the microvasculature early on.

Our study is limited by the lack of human tissue material before KF, and therefore, our study cannot demonstrate structural vascular changes in humans early on or molecular congruence between animals and humans. A limited number of samples and bulk tissue analyses preclude any definitive conclusions concerning the type of cell involvement or differences between the sexes. In addition, this study is limited by its relatively small study population and cross-sectional nature. Therefore, we cannot assess the predictive value of GABA, Hcy, and ADMA levels on disease progression.

In summary, our study demonstrates that in ADPKD, intrarenal MV abnormalities present early on and progress through the course of the disease. Importantly, intrarenal MV changes are preceded by both vascular transcriptional and metabolic alterations. Our data support that intrarenal microvessels may serve as a novel therapeutic target in ADPKD, and its underlying biomarkers may serve to monitor its progression.

## AUTHORS CONTRIBUTIONS

GY, SKS and BS performed experiments, data acquisition, and/or analysis. AA assisted with some data acquisition. WE, XX and YZ performed some experiments. CH, MCH and ARC provided insights into research findings. AE assisted with and supervised some data acquisition and analysis and provided insights into research findings. MVI designed and supervised the study, performed part of the data analysis, interpreted the results and research findings, and prepared the manuscript. All authors contributed to the review and edited the manuscript.

## ACKNOWLEDGMENTS

We acknowledge the Mayo Clinic Genome Analysis Core, Microscopy, Cell Analysis Cores, and the Mayo Clinic Translational PKD Center (MTPC). Special thanks to Prasanna Mishra and Ryan Meloche for continuously optimizing the NMR instruments when our experiments were performed.

## DATA SHARING STATEMENT

The active URL links to the datasets used in this manuscript are provided below: Vasculature-related genes at 1 month (Supplemental Data File 1) https://doi.org/10.6084/m9.figshare.28719626

Vasculature-related genes at 6 months (Supplemental Data File 2) https://doi.org/10.6084/m9.figshare.28719632

Vasculature-related genes at 12 months (Supplemental Data File 3) https://doi.org/10.6084/m9.figshare.28719638

## SUPPLEMENTAL MATERIAL

This article contains the following supplemental material online:

Supplemental Material (Detailed expanded methods, tables, figures and figure legends): https://doi.org/10.6084/m9.figshare.28719581

