## Supplemental material for "Vascular transcriptional and metabolic changes precede progressive intra-renal microvascular rarefaction in autosomal dominant polycystic kidney disease"

Running title: Renal microvasculature in ADPKD

Gizem Yilmaz[1,2*]; Santu K. Singha[1,2*]; Bansi Savaliya[1,2*]; Ahmed Abdelfattah[1,2]; Walaa Elsekaily[1,2]; Xiaohong Xu[1,2,3,4]; Youwen Zhang[1,2]; Christian Hanna[1,2,5]; Marie C. Hogan[1,2]; Alejandro R. Chade[6]; Alfonso Eirin[1,7]; and Maria V. Irazabal[1,2§]

[1]Department of Medicine, Division of Nephrology and Hypertension, Mayo Clinic, Rochester, MN

[2]Mayo Translational PKD Center, Mayo Clinic, Rochester, MN

[3]Department of Nephrology, The Affiliated Suqian First People's Hospital of Nanjing Medical University, Suqian, China;

[4]Department of Nephrology, Jiangsu Province Suqian Hospital, Suqian, China.

[5]Department of Pediatric and Adolescent Medicine, Division of Pediatric Nephrology and Hypertension, Mayo Clinic, Rochester, MN

[6]Department of Medical Pharmacology and Physiology, University of Missouri-Columbia, Missouri, MO

[7]Department of Cardiovascular Medicine, Mayo Clinic, Rochester, MN

*equally contributing

#### EXTENDED METHODS

*Animals study cohort, experimental design, and tissue harvest*

The study was approved by the Institutional Animal Care and Use Committee (IACUC, protocol #: A00003777 and A00005742). Animal experiments were performed to conform to NIH, USDA, and AALAS guidelines. All methods are reported in accordance with ARRIVE guidelines[1].

For this study, we used the *Pkd1*$^{RC/RC}$ mouse (a homozygous *Pkd1* model with the hypomorphic, human missense mutation p.Arg3277Cys), which is an established ADPKD model that reflects the slowly progressive nature of human disease [2]. *Pkd1*$^{RC/RC}$ animals were inbred into a mixed background (F1 progeny of 129S6/SvEvTac (129) *Pkd1*$^{RC/RC}$ and C57BL/6 (B6) *Pkd1*$^{RC/RC}$), and WT animals were in the same mixed background. *-8* No animals were lost during the study, and all were available for analysis. Experimenters were not blinded to the animal's genotype for morphological analyses because *Pkd1*$^{RC/RC}$ mice kidneys were visibly different from controls.

All animals were scanned by magnetic resonance imaging (MRI) at each time point, and TKV was determined. Systolic (SBP), diastolic (DBP), and mean arterial (MAP) pressure was monitored noninvasively using the tail-cuff method (CODA systems, Kent Scientific), and 24-hour urine samples were collected using metabolic cages. Animals were then returned to regular housing for 48 hours to minimize stress before euthanasia.

At the time of euthanasia, the left kidney was collected using our modified clamp-freezing protocol[3], a cardiac puncture was performed, blood was collected, and the right kidney was harvested after animal exsanguination in 10 animals (5 males and 5 females) per group. Plasma was used to measure blood urea nitrogen (BUN) using a colorimetric BUN assay (BUN-Urea, BioAssay Systems, Hayward, CA). The left kidney was pulverized in a liquid nitrogen-cooled mortar and kept at -80°C until sample preparation for mRNA-seq or quantitative polymerase chain reaction (qPCR). The right kidney was placed into vials containing 10% formaldehyde in phosphate buffer (pH 7.4), and tissues were embedded in paraffin for histology. Bodyweight (BW) and kidney weights were recorded after freezing (left) and before fixation (right). In randomly selected WT and *Pkd1*$^{RC/RC}$ mice (n=5, each per time point, 30 animals), RNA was extracted from homogenized whole left kidney tissue for mRNA-seq. In the remaining WT and *Pkd1*$^{RC/RC}$ mice (n=6, 3 males and 3 females at each per time point, 36 animals), animals were perfused with Microfil after euthanasia, and both right and left kidneys were prepared for 3D micro-CT studies.

*In vivo MRI analysis*

MRI scans were performed in a 16.4T Bruker Advance 700 MHz vertical bore nuclear magnetic resonance spectrometer using a 38 mm volume RF coil, as previously described[4]. Mouse temperature was maintained stable (35-37°C) throughout the scans. Respiratory gating was used, and images were acquired using a turbo Rapid Acquisition with Relaxation Enhancement (RARE) sequence, 11-19 coronal and axial slices with TR/TE 1,500/9 msec, RARE factor 8, (matrix 256 x 256, FOV 2.56 x 2.56 cm, slice thickness 0.75 mm). TKV was calculated from coronal slices using Analyze™ 12.0 software package (Biomedical Imaging Resource, Mayo Clinic, MN) and adjusted by animals' body length (blTKV).

*Histomorphometric Analysis*

Cystic area (CA) and fibrotic area (FA) were determined from 5µm paraffin-embedded longitudinal kidney tissue sections stained with hematoxylin-eosin (H&E) and Picosirus Red, respectively[5]. CA was calculated as the percentage of the total area of two coronally cut sections using the NIS elements software (Nikon Instruments Inc., Melville, NY). FA was calculated as the percentage of the total captured region from cortical and medullary images (n=4 each) and further adjusted by CA using MetaMorph software (Molecular Devices, Inc., San Jose, CA).

*Renal MV density, vessel diameter, and tortuosity*

The renal MV architecture was assessed using 3D micro-CT in WT and *Pkd1*$^{RC/RC}$ mice (n=6, 3 males and 3 females, 12 kidneys at each per time point, 36 animals). At the time of euthanasia, animals were weighed and anesthetized. Under a surgical level of anesthesia, a laparotomy (midline incision) through all layers up to the chest was performed after skin preparation (including clipping followed by the application of alcohol). The left ventricle was then cannulated with a 25-gauge butterfly infusion set, and a drop of vetbond was put on the puncture site to hold the needle in place. Blood was collected through the cardiac puncture, and an infusion of 0.9% saline (containing 10 units ⁄ mL heparin) was initiated under physiological perfusion pressure (Syringe Infusion Pump 22; Harvard Apparatus, Holliston, Massachusetts, USA)[6] at a rate of 2 mL⁄min. The right atrium was cut to allow the saline solution to drain. After 10–15 minutes of saline infusion and when it drained freely and clear from the right atrium, the saline syringe was replaced with a 20 mL syringe containing a freshly mixed radio-opaque silicone polymer (Microfil MV122; Flow Tech, Inc., Carver, Massachusetts, USA). The perfusion was continued with the contrast polymer agent through the same tubing at 1ml/min until the polymer drained freely from the atrium and all vessels were filled[7,8]. After the infusion was completed, the renal vessels were tied securely with a silk ligature, and both kidneys were left in place. The polymer-filled kidneys were allowed to cure at 4ºC for 24 hours before being placed in 10% formalin for at least 72 hours before scanning[9]. The microfil was diluted at 1:1:0.1, microfil, diluent, curing agent.

Intra-renal MV spatial density was calculated in a blinded manner by semi-automatically counting microvessels using the Analyze tool "Object Counter." Kidneys were tomographically divided into 15 levels (slices) from top to bottom, following the z-axis at equal intervals. A threshold for intensity was manually determined by observing the radio-opaque contrast (Microfil) in circumferential structures (vessels) of different diameters. Using the "Object Counter" tool, we calculated the number of renal microvessels of diameters between 0 and 500μm in each of the 15 2D z-axis sections and expressed as spatial density (number of vessels per tissue area, correcting for cystic area from MRI images). Circularity and rectangularity limits were used to exclude elliptical vessels that were not truly cross-sectional. The diameter of microvessels was also calculated in each cross-section. The cortex and medulla were divided into outer and inner regions[10] (OC, IC, OM, and IM), and MV density and vessel diameter were calculated at each level.

Tortuosity, an index of neovascularization and MV immaturity[11], was calculated in tomographically isolated renal microvessels, as described previously[12]. The entire 3D vascular path length (actual length) and linear length (the shortest distance from base to tip) were determined from the base of the main vessel at the cortico-medullary junction to its tip at the superficial cortex of 5 randomly selected vessels per kidney using the "Generate Tree Analysis" tool of Analyze. The elongation factor (tortuosity index) was obtained by dividing the path length by the linear length, and the results from all vessels were averaged.

*Peritubular capillary density*

Peritubular capillary density was determined in H&E-stained kidney sections (N=10 per animal) at x40 magnification (Carl ZEISS SMT, Oberkochen, Germany). Capillaries were identified by the presence of lumen, red blood cells and/or an endothelial cell lining[13]. The number of capillaries per field was calculated and further adjusted by CA, and expressed as the number of capillaries/parenchyma[5].

*Perivascular fibrosis*

Perivascular fibrosis was determined in Masson's trichrome-stained slides at x20 magnification. The percentage of fibrosis was quantified in 15 random similar-sized (major diameter ranging from 0-1000μm) renal vessels per animal using the NIH software Image J and expressed as % fibrosis/parenchyma.

*mRNA-seq and Bioinformatic Analysis*

mRNA-seq was performed at the Mayo Clinic Genomic Analysis Core, as previously described[14,15]. The total RNA was extracted at the Mayo Clinic Biospecimens Accessioning and Processing Core (BAP) using Qiagen QIAcube according to the manufacturer's instructions. After mRNA extraction, QC was performed before library preparation. All samples passed quality control and obtained RNA integrity numbers >7.0 as recommended for the library preparation. RNA libraries were prepared using TruSeq RNA Sample Prep Kit v2, and samples were sequenced on an Illumina HiSeq 2000 using TruSeq SBS sequencing kit version 3 and HCS v2.0.12 data collection software. Paired-end reads were processed using our RNA-seq bioinformatics pipeline (MAP-RSeq version 3.1.4)[16]. Reads were aligned using the MAP-RSeq tool STAR[17], aligned reads were further processed, and gene and exon expression was quantified using the Subread aligner[18]. Gene expression data were normalized by Counts per Million (CPM). Single nucleotide variants (SNVs), small insertions-deletions (indels), as well as known and novel gene isoforms were detected using[17], GATK[19], Haplotype caller[19], RVBoost[20], and StringTie[21]. Indepth analyses were performed on aligned reads to examine the quality of the sequenced libraries, and differential exon use was assessed using DEXSeq[22], before reporting results by MAP-RSeq. To visualize the whole spectrum of changes in the expression of vasculature-related genes in WT and *Pkd1*$^{RC/RC}$ mouse kidneys at 1, 6, and 12 months, we used the Mouse Genome Informatics (MGI) database to screen genes associated with angiogenesis (MGI_20240205_142825).

Differential expression analysis of vasculature-related genes was performed using raw gene counts from MAP-RSeq. Differentially expressed genes (DEGs) (CPM>0.1, fold-change >1.4 or <0.7, and p values <0.05) were identified using edgeR[23], FDR-corrected using the Benjamini-Hochberg-Yekutieli procedure[24], reported along with their magnitude of change, and visualized in volcano plots (Excel, Microsoft) and heat maps (Heatmapper, www.heatmappear.ca). (**Supp. Data files 1-3**)

DEGs were further classified by their cellular component, molecular function, and biological process using Gene Set Enrichment Analysis (GSEA)[25], and interrogation of protein functional and physical interactions was performed using the Search Tool for the Retrieval of Interacting Genes (STRING) v9.1 (http://string-db.org/)[26].

*Validation of mRNA-seq analysis*

Validation of mRNA-seq was performed in randomly selected top-upregulated (FC >2.5, p<0.001) and top-downregulated (FC <0.4, p<0.001) mRNAs in *Pkd1*$^{RC/RC}$ compared to WT kidneys at 1, 6 and 12 months time points. *Wnt7a* and *Sfrp2* (1 month), *Ccl12* and *Sfrp2* (6 months), and *Wnt7a* and *Adtrp* (12 months) were chosen for validation, and their expression levels were determined by real-time qPCR using the ΔΔCt method[27]. All primers were obtained from Life Technologies Corporation (Carlsbad, CA, assay ID: 00437356, 01213947, 01617100, and 01336647). Kidney tissue lysate was processed to isolate cDNA according to manufacturer guidelines (PureLink RNA Mini Kit, Thermo Fisher Scientific, Rochester, MN). mRNA expression was normalized to β-actin (*Actb*), and results were expressed relative to WT controls.

*Human subject cohort*

The study was approved by our Institutional Review Board (IRB) of the Mayo Clinic in accordance with the Declaration of Helsinki and the Health Insurance Portability and Accountability Act (HIPAA) guidelines and conducted under IRB 24-013591. Thirty-two, young individuals (8-35 years of age), with SBP <130 mmHg) without antihypertensive medication (in individuals >13 years of age) or <95th%ile for height, age and gender (in individuals <13 years of age)[28], with eGFR>90mL/min/1.73m², and with a previous diagnosis of ADPKD based on Ravine/Pei criteria[29,30] were matched 2:1 for age (±2 years) and gender to control individuals without a personal or family history of kidney disease and included in this cross-sectional study. Exclusion criteria included patients with atypical presentation of the disease as described[31], antihypertensive medication, diabetes mellitus, predicted urine protein excretion >1g/24hrs, abnormal urinalysis suggestive of concomitant glomerular disease, and concomitant systemic disease known to affect the kidney (e.g. lupus, hepatitis B or C, amyloidosis), or a known pathogenic variant in other than in *PKD1* or *PKD2* genes.

On the day of the study, subjects completed a health and family questionnaire and collected 2nd-morning urine after a fasting period of 10 hours for a urinalysis and, if applicable, exclusion of pregnancy and a blood sample for chemistries. An abdominal MRI without gadolinium was acquired on a GE 3T scanner (GE Medical Systems, Discovery MR750w) for the estimation of TKV (in n=28 individuals with ADPKD and n=14 controls) and RBF (in n=21 individuals with ADPKD and n=14 controls), as previously described[32-34]. TKVs were adjusted by the patient's height (htTKV) and expressed as ml/m, and RBF was adjusted by the patient's body surface area (BSA) and expressed as mL/min/1.73m$^2$. Patients in which htTKV was available (n=28), were classified based on their htTKV and age as described[31]. Blood pressure, height, and weight were obtained at the time of the MRI. The eGFR was calculated with the Chronic Kidney Disease in Children (CKiD) equation (revised/bedside Schwartz equation) for individuals <18 years old and with the Chronic Kidney Disease Epidemiology Collaboration equation (2021) for individuals >18 years[35,36]. Genetic data was available in 19 individuals with ADPKD.

*Pilot metabolomics analyses*

The concentrations of several metabolites involved in cellular energy metabolism including the tricarboxylic acid (TCA) cycle analytes, the glycolytic pathway, fatty acids (FAs) and amino acids (AAs) were determined by liquid chromatography-tandem mass spectrometry (LC-MS/MS) metabolomic analysis (n=10 individuals with ADPKD and 10 matched controls). Plasma samples were aliquoted and frozen within 30 minutes of collection. All samples were stored at -80°C until completion of the study. Plasma metabolite levels were expressed as µM. TCA analytes were measured by gas chromatograph mass spectrometry (GC/MS) as previously described[37-39] with a few modifications. Briefly, 50ul of plasma were mixed with 20ul of internal solution containing U-$^{13}$C labeled analytes. The proteins were removed by adding 300 ul of chilled methanol and acetonitrile solution to the sample mixture. After drying the supernatant in the speed vac, the sample was derivatized with ethoxime and then with MtBSTFA + 1% tBDMCS (N-Methyl-N-(t-Butyldimethylsilyl)-Trifluoroacetamide + 1% t-Butyldimethylchlorosilane) before it was analyzed on an Agilent 5977B GC/MS (gas chromatography/mass spectrometry) under electron impact and single ion monitoring conditions. Concentrations of lactic acid (m/z 261.2), fumaric acid (m/z 287.1), succinic acid (m/z 289.1), ketoglutaric acid (m/z 360.2), malic acid (m/z 419.3), aspartic acid (m/z 418.2), citric acid (m/z 591.4), and glutamic acid (m/z 432.4) were measured against 7-point calibration curves that underwent the same derivatization. FAs concentrations were measured against a standard curve on a triple quadrupole mass spectrometer coupled with an Ultra Pressure Liquid Chromatography system (LC/MS) as previously described[40]. AAs and their metabolites were measured by LCMS as previously described[41,42]. Data was acquired under negative electrospray ionization conditions in selective reaction monitoring mode. FAs and AAs concentrations were established by comparing their ion intensity (121-labeled AAs and FAs) to their respective internal standards (113-labeled AAs and FAs)[43].

*Gene–metabolite interaction analysis*

Gene–metabolite interaction networks were generated using MetaboAnalyst v.5.0 (University of Alberta RRID:SCR_015539) by mapping vasculature-related DEGs in *Pkd1*$^{RC/RC}$ at 1 month and metabolites dysregulated in individuals with ADPKD using a comprehensive gene-metabolite interaction data on STITCH ('search tool for interactions of chemicals')[44].

*Confirmatory studies in plasma samples from individuals with ADPKD and controls*

Plasma Hcy, gamma-aminobutyric acid (GABA) and ADMA levels were measured by enzyme-linked immunosorbent assay (ELISA, Cell Biolabs, Cat. no. STA-670, LifeSpan BioSciences, Cat. no. LS-F10676 and Eagle Biosciences, Cat. No. ADM31-K01), in n=32 individuals with ADPKD and 16 age and sex-matched controls, according to manufacturer protocols and expressed as ng/mL.

*Statistical analysis*

Statistical analysis of animal clinical and laboratory characteristics, as well as patient clinical and laboratory characteristics, were performed using PRISM9 (GraphPad Software, La Jolla, CA). The statistical analysis of metabolomics data was conducted using MetaboAnalyst v.5.0 (University of Alberta RRID:SCR_015539) after performing auto-scaling, which uses each variable's standard deviation as the scaling factor[45]. The Shapiro-Wilk test was used to test for deviation from normality. Normally distributed data were expressed as means ± SD and compared using parametric (Student's t-test) methods. In contrast, data that did not follow a Gaussian distribution were expressed as medians (interquartile ranges) and compared using nonparametric (Kruskal-Wallis) tests. Regressions were calculated by the least-squares fit (parametric) and Spearman rank correlation coefficient (nonparametric). Longitudinal data were analyzed with parametric (paired t test) and nonparametric (Wilcoxon Signed-Rank) tests. p values ≤0.05 were considered significant.

#### SUPPLEMENTAL TABLES

**Table S1. Vasculature related genes**

| ***Symbol*** | ***Gene Name*** |
|---|---|
| Abcc8 | ATP-binding cassette, sub-family C member 8 |
| Adamts1 | ADAM metallopeptidase with thrombospondin type 1 motif 1 |
| Adrb2 | adrenergic receptor, beta 2 |
| Ago1 | argonaute RISC catalytic subunit 1 |
| Agt | angiotensinogen |
| Alox5 | arachidonate 5-lipoxygenase |
| Apoh | apolipoprotein H |
| Atf2 | activating transcription factor 2 |
| Atp2b4 | ATPase, Ca++ transporting, plasma membrane 4 |
| Ccn6 | cellular communication network factor 6 |
| Cd59a | CD59a antigen |
| Cd160 | CD160 antigen |
| Cldn5 | claudin 5 |
| Cnmd | chondromodulin |
| Col4a3 | collagen, type IV, alpha 3 |
| Crhr2 | corticotropin releasing hormone receptor 2 |
| Ctnnb1 | catenin beta 1 |
| Cx3cr1 | C-X3-C motif chemokine receptor 1 |
| Cxcl10 | C-X-C motif chemokine ligand 10 |
| Dcn | decorin |
| Efna3 | ephrin A3 |
| Emilin1 | elastin microfibril interfacer 1 |
| Fasl | Fas ligand |
| Fbln5 | fibulin 5 |
| Flcn | folliculin |
| Foxj2 | forkhead box J2 |
| Foxo4 | forkhead box O4 |
| Gadd45a | growth arrest and DNA-damage-inducible 45 alpha |
| Gtf2i | general transcription factor II I |
| Hgs | HGF-regulated tyrosine kinase substrate |
| Hhex | hematopoietically expressed homeobox |
| Hhip | Hedgehog-interacting protein |
| Hoxa5 | homeobox A5 |
| Il17f | interleukin 17F |
| Klf4 | Kruppel-like transcription factor 4 (gut) |
| Lif | leukemia inhibitory factor |
| Mecp2 | methyl CpG binding protein 2 |
| Mir329 | microRNA 329 |

| | |
|---|---|
| Naxe | NAD(P)HX epimerase |
| Ngfr | nerve growth factor receptor (TNFR superfamily, member 16) |
| Ngp | neutrophilic granule protein |
| Pgk1 | phosphoglycerate kinase 1 |
| Plg | plasminogen |
| Plk2 | polo like kinase 2 |
| Pml | promyelocytic leukemia |
| Pparg | peroxisome proliferator activated receptor gamma |
| Prl7d1 | prolactin family 7, subfamily d, member 1 |
| Ptn | pleiotrophin |
| Ptprm | protein tyrosine phosphatase receptor type M |
| Qki | quaking, KH domain containing RNA binding |
| Rela | v-rel reticuloendotheliosis viral oncogene homolog A (avian) |
| Rgcc | regulator of cell cycle |
| Rock1 | Rho-associated coiled-coil containing protein kinase 1 |
| Rock2 | Rho-associated coiled-coil containing protein kinase 2 |
| S2bpcox16 | synaptojanin 2 binding protein Cox16 readthrough |
| Sars1 | seryl-tRNA synthetase 1 |
| Serpinf1 | serine (or cysteine) peptidase inhibitor, clade F, member 1 |
| Sparc | secreted acidic cysteine rich glycoprotein |
| Spred1 | sprouty protein with EVH-1 domain 1, related sequence |
| Spry2 | sprouty RTK signaling antagonist 2 |
| Stab1 | stabilin 1 |
| Stat1 | signal transducer and activator of transcription 1 |
| Sulf1 | sulfatase 1 |
| Synj2bp | synaptojanin 2 binding protein |
| Tcf4 | transcription factor 4 |
| Tgfb2 | transforming growth factor, beta 2 |
| Thbs2 | thrombospondin 2 |
| Tnmd | tenomodulin |
| Il12a | interleukin 12a |
| Il12b | interleukin 12b |
| Sema6a | sema domain, transmembrane domain (TM), and cytoplasmic domain, (semaphorin) 6A |
| Card10 | caspase recruitment domain family, member 10 |
| Hdac5 | histone deacetylase 5 |
| Map2k5 | mitogen-activated protein kinase kinase 5 |
| Pik3r2 | phosphoinositide-3-kinase regulatory subunit 2 |
| Rhoa | ras homolog family member A |
| Adamts9 | ADAM metallopeptidase with thrombospondin type 1 motif 9 |
| Creb3l1 | cAMP responsive element binding protein 3-like 1 |
| E2f2 | E2F transcription factor 2 |
| Epn1 | epsin 1 |

| | |
|---|---|
| Epn2 | epsin 2 |
| Klf2 | Kruppel-like transcription factor 2 (lung) |
| Sh2b3 | SH2B adaptor protein 3 |
| Stard13 | StAR related lipid transfer domain containing 13 |
| Slc12a2 | solute carrier family 12, member 2 |
| Tafa5 | TAFA chemokine like family member 5 |
| Tnf | tumor necrosis factor |
| Ackr3 | atypical chemokine receptor 3 |
| Actg1 | actin, gamma, cytoplasmic 1 |
| Acvrl1 | activin A receptor, type II-like 1 |
| Adam8 | a disintegrin and metallopeptidase domain 8 |
| Adam15 | ADAM metallopeptidase domain 15 |
| Adgra2 | adhesion G protein-coupled receptor A2 |
| Adgrg1 | adhesion G protein-coupled receptor G1 |
| Adm2 | adrenomedullin 2 |
| Adra2b | adrenergic receptor, alpha 2b |
| Aggf1 | angiogenic factor with G patch and FHA domains 1 |
| Aimp1 | aminoacyl tRNA synthetase complex-interacting multifunctional protein 1 |
| Amot | angiomotin |
| Amotl1 | angiomotin-like 1 |
| Amotl2 | angiomotin-like 2 |
| Ang | angiogenin, ribonuclease, RNase A family, 5 |
| Ang2 | angiogenin, ribonuclease A family, member 2 |
| Ang3 | angiogenin, ribonuclease A family, member 3 |
| Ang4 | angiogenin, ribonuclease A family, member 4 |
| Ang5 | angiogenin, ribonuclease A family, member 5 |
| Ang6 | angiogenin, ribonuclease A family, member 6 |
| Angpt1 | angiopoietin 1 |
| Angpt2 | angiopoietin 2 |
| Angpt4 | angiopoietin 4 |
| Angptl3 | angiopoietin-like 3 |
| Angptl4 | angiopoietin-like 4 |
| Angptl6 | angiopoietin-like 6 |
| Anpep | alanyl aminopeptidase, membrane |
| Anxa2 | annexin A2 |
| Apela | apelin receptor early endogenous ligand |
| Apln | apelin |
| Aplnr | apelin receptor |
| Apold1 | apolipoprotein L domain containing 1 |
| Arhgap22 | Rho GTPase activating protein 22 |
| Arhgap24 | Rho GTPase activating protein 24 |
| Atp5f1b | ATP synthase F1 subunit beta |
| Bcas3 | BCAS3 microtubule associated cell migration factor |

| | |
|---|---|
| Becn1 | beclin 1, autophagy related |
| Bmp4 | bone morphogenetic protein 4 |
| Bmpr1a | bone morphogenetic protein receptor, type 1A |
| Bsg | basigin |
| C1galt1 | core 1 synthase, glycoprotein-N-acetylgalactosamine 3-beta-galactosyltransferase, 1 |
| Calcrl | calcitonin receptor-like |
| Cald1 | caldesmon 1 |
| Casp8 | caspase 8 |
| Cav1 | caveolin 1, caveolae protein |
| Ccbe1 | collagen and calcium binding EGF domains 1 |
| Ccdc134 | coiled-coil domain containing 134 |
| Ccl2 | C-C motif chemokine ligand 2 |
| Ccl12 | C-C motif chemokine ligand 12 |
| Ccn2 | cellular communication network factor 2 |
| Ccn3 | cellular communication network factor 3 |
| Ccr2 | C-C motif chemokine receptor 2 |
| Cd47 | CD47 antigen (Rh-related antigen, integrin-associated signal transducer) |
| Cemip2 | cell migration inducing hyaluronidase 2 |
| Cfh | complement component factor h |
| Cib1 | calcium and integrin binding 1 |
| Clic4 | chloride intracellular channel 4 |
| Col4a1 | collagen, type IV, alpha 1 |
| Col4a2 | collagen, type IV, alpha 2 |
| Col8a1 | collagen, type VIII, alpha 1 |
| Col8a2 | collagen, type VIII, alpha 2 |
| Col18a1 | collagen, type XVIII, alpha 1 |
| Col27a1 | collagen, type XXVII, alpha 1 |
| Cspg4 | chondroitin sulfate proteoglycan 4 |
| Cxcl17 | C-X-C motif chemokine ligand 17 |
| Cxcr3 | C-X-C motif chemokine receptor 3 |
| Cyp1b1 | cytochrome P450, family 1, subfamily b, polypeptide 1 |
| Dab2ip | disabled 2 interacting protein |
| Dicer1 | dicer 1, ribonuclease type III |
| Dll4 | delta like canonical Notch ligand 4 |
| Dysf | dysferlin |
| Ecm1 | extracellular matrix protein 1 |
| Ecscr | endothelial cell surface expressed chemotaxis and apoptosis regulator |
| Edn2 | endothelin 2 |
| Ednra | endothelin receptor type A |
| Efna1 | ephrin A1 |
| Efnb2 | ephrin B2 |
| Egf | epidermal growth factor |
| Egfl7 | EGF-like domain 7 |

| | |
|---|---|
| Eif2ak3 | eukaryotic translation initiation factor 2 alpha kinase 3 |
| Elk3 | ELK3, member of ETS oncogene family |
| Emc10 | ER membrane protein complex subunit 10 |
| Eng | endoglin |
| Enpep | glutamyl aminopeptidase |
| Epas1 | endothelial PAS domain protein 1 |
| Epgn | epithelial mitogen |
| Epha1 | Eph receptor A1 |
| Epha2 | Eph receptor A2 |
| Epha5 | Eph receptor A5 |
| Ephb1 | Eph receptor B1 |
| Ephb2 | Eph receptor B2 |
| Ephb3 | Eph receptor B3 |
| Ephb4 | Eph receptor B4 |
| Epo | erythropoietin |
| Ereg | epiregulin |
| Esm1 | endothelial cell-specific molecule 1 |
| Fap | fibroblast activation protein |
| Fgf1 | fibroblast growth factor 1 |
| Fgf2 | fibroblast growth factor 2 |
| Fgf6 | fibroblast growth factor 6 |
| Fgf9 | fibroblast growth factor 9 |
| Fgf18 | fibroblast growth factor 18 |
| Fgfr1 | fibroblast growth factor receptor 1 |
| Fgfr2 | fibroblast growth factor receptor 2 |
| Flna | filamin, alpha |
| Flt1 | FMS-like tyrosine kinase 1 |
| Flt4 | FMS-like tyrosine kinase 4 |
| Fmnl3 | formin-like 3 |
| Fn1 | fibronectin 1 |
| Foxc1 | forkhead box C1 |
| Fzd4 | frizzled class receptor 4 |
| Fzd5 | frizzled class receptor 5 |
| Fzd8 | frizzled class receptor 8 |
| Gab1 | growth factor receptor bound protein 2-associated protein 1 |
| Gata2 | GATA binding protein 2 |
| Gdf2 | growth differentiation factor 2 |
| Glul | glutamate-ammonia ligase |
| Gna13 | guanine nucleotide binding protein, alpha 13 |
| Gpr15 | G protein-coupled receptor 15 |
| Grem1 | gremlin 1, DAN family BMP antagonist |
| Hand1 | heart and neural crest derivatives expressed 1 |
| Hand2 | heart and neural crest derivatives expressed 2 |

| | |
|---|---|
| Hbegf | heparin-binding EGF-like growth factor |
| Hif1a | hypoxia inducible factor 1, alpha subunit |
| Hif3a | hypoxia inducible factor 3, alpha subunit |
| Hmox1 | heme oxygenase 1 |
| Hrg | histidine-rich glycoprotein |
| Hs6st1 | heparan sulfate 6-O-sulfotransferase 1 |
| Hspg2 | perlecan (heparan sulfate proteoglycan 2) |
| Htatip2 | HIV-1 Tat interactive protein 2 |
| Il18 | interleukin 18 |
| Itga2b | integrin alpha 2b |
| Itga5 | integrin alpha 5 (fibronectin receptor alpha) |
| Itgav | integrin alpha V |
| Itgb1bp1 | integrin beta 1 binding protein 1 |
| Jam3 | junction adhesion molecule 3 |
| Jun | jun proto-oncogene |
| Kctd10 | potassium channel tetramerisation domain containing 10 |
| Kdr | kinase insert domain protein receptor |
| Klf5 | Kruppel-like transcription factor 5 |
| Krit1 | KRIT1, ankyrin repeat containing |
| Lemd3 | LEM domain containing 3 |
| Lep | leptin |
| Lepr | leptin receptor |
| Map3k7 | mitogen-activated protein kinase kinase kinase 7 |
| Mapk14 | mitogen-activated protein kinase 14 |
| Mcam | melanoma cell adhesion molecule |
| Med1 | mediator complex subunit 1 |
| Meis1 | Meis homeobox 1 |
| Meox2 | mesenchyme homeobox 2 |
| Mfge8 | milk fat globule EGF and factor V/VIII domain containing |
| Minar1 | membrane integral NOTCH2 associated receptor 1 |
| Minar2 | membrane integral NOTCH2 associated receptor 2 |
| Mir126b | microRNA 126b |
| Mmp2 | matrix metallopeptidase 2 |
| Mmp19 | matrix metallopeptidase 19 |
| Mmrn2 | multimerin 2 |
| Mydgf | myeloid derived growth factor |
| Myh9 | myosin, heavy polypeptide 9, non-muscle |
| Naa15 | N(alpha)-acetyltransferase 15, NatA auxiliary subunit |
| Ncl | nucleolin |
| Ndnf | neuron-derived neurotrophic factor |
| Ndp | Norrie disease (pseudoglioma) (human) |
| Nf1 | neurofibromin 1 |
| Ninj1 | ninjurin 1 |

| | |
|---|---|
| Nos3 | nitric oxide synthase 3, endothelial cell |
| Notch1 | notch 1 |
| Nox1 | NADPH oxidase 1 |
| Nppc | natriuretic peptide type C |
| Npr3 | natriuretic peptide receptor 3 |
| Nr2e1 | nuclear receptor subfamily 2, group E, member 1 |
| Nrp1 | neuropilin 1 |
| Nrp2 | neuropilin 2 |
| Nrxn1 | neurexin I |
| Nrxn3 | neurexin III |
| Nus1 | NUS1 dehydrodolichyl diphosphate synthase subunit |
| Optc | opticin |
| Or10j5 | olfactory receptor family 10 subfamily J member 5 |
| Otulin | OTU deubiquitinase with linear linkage specificity |
| Ovol2 | ovo like zinc finger 2 |
| Pank2 | pantothenate kinase 2 |
| Parva | parvin, alpha |
| Pdcd6 | programmed cell death 6 |
| Pdcd10 | programmed cell death 10 |
| Pdcl3 | phosducin-like 3 |
| Pde3b | phosphodiesterase 3B, cGMP-inhibited |
| Pdgfa | platelet derived growth factor, alpha |
| Pdgfrb | platelet derived growth factor receptor, beta polypeptide |
| Pecam1 | platelet/endothelial cell adhesion molecule 1 |
| Pgf | placental growth factor |
| Pik3ca | phosphatidylinositol-4,5-bisphosphate 3-kinase catalytic subunit alpha |
| Pik3cg | phosphatidylinositol-4,5-bisphosphate 3-kinase catalytic subunit gamma |
| Pik3r6 | phosphoinositide-3-kinase regulatory subunit 5 |
| Pknox1 | Pbx/knotted 1 homeobox |
| Plau | plasminogen activator, urokinase |
| Plcd1 | phospholipase C, delta 1 |
| Plcd3 | phospholipase C, delta 3 |
| Plxnd1 | plexin D1 |
| Pnpla6 | patatin-like phospholipase domain containing 6 |
| Pofut1 | protein O-fucosyltransferase 1 |
| Prkca | protein kinase C, alpha |
| Prkd1 | protein kinase D1 |
| Prkd2 | protein kinase D2 |
| Prkx | protein kinase, X-linked |
| Prok1 | prokineticin 1 |
| Prok2 | prokineticin 2 |
| Psg22 | pregnancy-specific beta-1-glycoprotein 22 |
| Pten | phosphatase and tensin homolog |

| | |
|---|---|
| Ptgs2 | prostaglandin-endoperoxide synthase 2 |
| Ptk2 | PTK2 protein tyrosine kinase 2 |
| Ptk2b | PTK2 protein tyrosine kinase 2 beta |
| Ptprb | protein tyrosine phosphatase receptor type B |
| Pxdn | peroxidasin |
| Ramp1 | receptor (calcitonin) activity modifying protein 1 |
| Ramp2 | receptor (calcitonin) activity modifying protein 2 |
| Rapgef3 | Rap guanine nucleotide exchange factor (GEF) 3 |
| Rasip1 | Ras interacting protein 1 |
| Rbpj | recombination signal binding protein for immunoglobulin kappa J region |
| Rhob | ras homolog family member B |
| Rhoj | ras homolog family member J |
| Rnf213 | ring finger protein 213 |
| Robo4 | roundabout guidance receptor 4 |
| Rora | RAR-related orphan receptor alpha |
| Rspo3 | R-spondin 3 |
| Rtl1 | retrotransposon Gaglike 1 |
| Rtn4 | reticulon 4 |
| S1pr1 | sphingosine-1-phosphate receptor 1 |
| Scg2 | secretogranin II |
| Sema3e | sema domain, immunoglobulin domain (Ig), short basic domain, secreted, (semaphorin) 3E |
| Sema4a | sema domain, immunoglobulin domain (Ig), transmembrane domain (TM) and short cytoplasmic domain, (semaphorin) 4A |
| Serpine1 | serine (or cysteine) peptidase inhibitor, clade E, member 1 |
| Setd2 | SET domain containing 2 |
| Shb | src homology 2 domain-containing transforming protein B |
| Shc1 | src homology 2 domain-containing transforming protein C1 |
| Shh | sonic hedgehog |
| Sirt1 | sirtuin 1 |
| Slc31a1 | solute carrier family 31, member 1 |
| Smad5 | SMAD family member 5 |
| Sox17 | SRY (sex determining region Y)-box 17 |
| Sox18 | SRY (sex determining region Y)-box 18 |
| Srpk2 | serine/arginine-rich protein specific kinase 2 |
| Srpx2 | sushi-repeat-containing protein, X-linked 2 |
| Syk | spleen tyrosine kinase |
| Tal1 | T cell acute lymphocytic leukemia 1 |
| Tbx1 | T-box 1 |
| Tbx4 | T-box 4 |
| Tek | TEK receptor tyrosine kinase |
| Tgfa | transforming growth factor alpha |
| Tgfbr1 | transforming growth factor, beta receptor I |
| Thbs1 | thrombospondin 1 |

| | |
|---|---|
| Thsd7a | thrombospondin, type I, domain containing 7A |
| Thy1 | thymus cell antigen 1, theta |
| Tie1 | tyrosine kinase with immunoglobulin-like and EGF-like domains 1 |
| Tmem100 | transmembrane protein 100 |
| Tmem215 | transmembrane protein 215 |
| Tnfaip2 | tumor necrosis factor, alpha-induced protein 2 |
| Tnfrsf12a | tumor necrosis factor receptor superfamily, member 12a |
| Tnfsf12 | tumor necrosis factor (ligand) superfamily, member 12 |
| Tspan12 | tetraspanin 12 |
| Ubp1 | upstream binding protein 1 |
| Unc5b | unc-5 netrin receptor B |
| Vash1 | vasohibin 1 |
| Vav2 | vav 2 oncogene |
| Vav3 | vav 3 oncogene |
| Vegfa | vascular endothelial growth factor A |
| Vegfb | vascular endothelial growth factor B |
| Vegfc | vascular endothelial growth factor C |
| Vegfd | vascular endothelial growth factor D |
| Vezf1 | vascular endothelial zinc finger 1 |
| Vhl | von Hippel-Lindau tumor suppressor |
| Vps4b | vacuolar protein sorting 4B |
| Wars1 | tryptophanyl-tRNA synthetase1 |
| Wasf2 | WASP family, member 2 |
| Wnt7a | wingless-type MMTV integration site family, member 7A |
| Wnt7b | wingless-type MMTV integration site family, member 7B |
| Xbp1 | X-box binding protein 1 |
| Ywhaz | tyrosine 3-monooxygenase/tryptophan 5-monooxygenase activation protein, zeta polypeptide |
| Zc3h12a | zinc finger CCCH type containing 12A |
| Angvq1 | angiogenesis by VEGF QTL 1 |
| Angvq2 | angiogenesis by VEGF QTL 2 |
| Angfq1 | angiogenesis due to FGF2 QTL 1 |
| Angfq2 | angiogenesis due to FGF2 QTL 2 |
| Angfq3 | angiogenesis due to FGF2 QTL 3 |
| Angfq4 | angiogenesis due to FGF2 QTL 4 |
| Angfq5 | angiogenesis due to FGF2 QTL 5 |
| Angfq7 | angiogenesis due to FGF2 QTL 7 |
| Angfq8 | angiogenesis due to FGF2 QTL 8 |
| Ace | angiotensin I converting enzyme |
| Rxra | retinoid X receptor alpha |
| B4galt1 | UDP-Gal:betaGlcNAc beta 1,4- galactosyltransferase, polypeptide 1 |
| Cx3cl1 | C-X3-C motif chemokine ligand 1 |
| Gpr4 | G protein-coupled receptor 4 |

| | |
|---|---|
| Gpx1 | glutathione peroxidase 1 |
| Hpse | heparanase |
| Pik3cb | phosphatidylinositol-4,5-bisphosphate 3-kinase catalytic subunit beta |
| Prcp | prolylcarboxypeptidase (angiotensinase C) |
| Nrarp | Notch-regulated ankyrin repeat protein |
| Baiap2 | brain-specific angiogenesis inhibitor 1-associated protein 2 |
| Baiap2l1 | BAI1-associated protein 2-like 1 |
| Baiap2l2 | BAI1-associated protein 2-like 2 |
| Acvr1 | activin A receptor, type 1 |
| Ahr | aryl-hydrocarbon receptor |
| Cxcl12 | C-X-C motif chemokine ligand 12 |
| Edn1 | endothelin 1 |
| Fgf8 | fibroblast growth factor 8 |
| Gbx2 | gastrulation brain homeobox 2 |
| Ihh | Indian hedgehog |
| Nfatc3 | nuclear factor of activated T cells, cytoplasmic, calcineurin dependent 3 |
| Nfatc4 | nuclear factor of activated T cells, cytoplasmic, calcineurin dependent 4 |
| Notch4 | notch 4 |
| Pitx2 | paired-like homeodomain transcription factor 2 |
| Ppp3r1 | protein phosphatase 3, regulatory subunit B, alpha isoform (calcineurin B, type I) |
| Rbm15 | RNA binding motif protein 15 |
| Sema5a | sema domain, seven thrombospondin repeats (type 1 and type 1-like), transmembrane domain (TM) and short cytoplasmic domain, (semaphorin) 5A |
| Stk4 | serine/threonine kinase 4 |
| Tbx20 | T-box 20 |
| Vangl2 | VANGL planar cell polarity 2 |
| Adtrp | androgen dependent TFPI regulating protein |
| Akt1 | thymoma viral proto-oncogene 1 |
| Egr3 | early growth response 3 |
| Gpld1 | glycosylphosphatidylinositol specific phospholipase D1 |
| Mia3 | MIA SH3 domain ER export factor 3 |
| Nr4a1 | nuclear receptor subfamily 4, group A, member 1 |
| Pik3r3 | phosphoinositide-3-kinase regulatory subunit 3 |
| Robo1 | roundabout guidance receptor 1 |
| Slit2 | slit guidance ligand 2 |
| Srf | serum response factor |
| Notch2 | notch 2 |
| Notch3 | notch 3 |
| Adgrb1 | adhesion G protein-coupled receptor B1 |
| Adgrb2 | adhesion G protein-coupled receptor B2 |
| Adgrb3 | adhesion G protein-coupled receptor B3 |
| Ccn1 | cellular communication network factor 1 |
| Ism1 | isthmin 1, angiogenesis inhibitor |

| | |
|---|---|
| Pdgfra | platelet derived growth factor receptor, alpha polypeptide |
| Tcf21 | transcription factor 21 |
| Adam12 | ADAM metallopeptidase domain 12 |
| Add1 | adducin 1 |
| Adm | adrenomedullin |
| Ago2 | argonaute RISC catalytic subunit 2 |
| Akt3 | thymoma viral proto-oncogene 3 |
| Anxa3 | annexin A3 |
| Aqp1 | aquaporin 1 |
| Brca1 | breast cancer 1, early onset |
| Btg1 | BTG anti-proliferation factor 1 |
| C3 | complement component 3 |
| C3ar1 | complement component 3a receptor 1 |
| C5ar1 | complement component 5a receptor 1 |
| C6 | complement component 6 |
| Camp | cathelicidin antimicrobial peptide |
| Ccl5 | C-C motif chemokine ligand 5 |
| Ccl11 | C-C motif chemokine ligand 11 |
| Ccl24 | C-C motif chemokine ligand 24 |
| Ccr3 | C-C motif chemokine receptor 3 |
| Cd34 | CD34 antigen |
| Cd40 | CD40 antigen |
| Cdh5 | cadherin 5 |
| Cela1 | chymotrypsin-like elastase family, member 1 |
| Chi3l1 | chitinase 3 like 1 |
| Cma1 | chymase 1, mast cell |
| Ctsh | cathepsin H |
| Cxcr2 | C-X-C motif chemokine receptor 2 |
| Cybb | cytochrome b-245, beta polypeptide |
| Cysltr1 | cysteinyl leukotriene receptor 1 |
| Cysltr2 | cysteinyl leukotriene receptor 2 |
| Ddah1 | dimethylarginine dimethylaminohydrolase 1 |
| Emilin2 | elastin microfibril interfacer 2 |
| Erap1 | endoplasmic reticulum aminopeptidase 1 |
| F3 | coagulation factor III |
| Gata4 | GATA binding protein 4 |
| Gata6 | GATA binding protein 6 |
| Grn | granulin |
| Hc | hemolytic complement |
| Hgf | hepatocyte growth factor |
| Hipk2 | homeodomain interacting protein kinase 2 |
| Hk2 | hexokinase 2 |
| Hmga2 | high mobility group AT-hook 2 |

| | |
|---|---|
| Hspb1 | heat shock protein 1 |
| Hspb6 | heat shock protein, alpha-crystallin-related, B6 |
| Hyal1 | hyaluronoglucosaminidase 1 |
| Igf2 | insulin-like growth factor 2 |
| Il1a | interleukin 1 alpha |
| Il1b | interleukin 1 beta |
| Isl1 | ISL1 transcription factor, LIM/homeodomain |
| Itgax | integrin alpha X |
| Itgb1 | integrin beta 1 (fibronectin receptor beta) |
| Itgb2 | integrin beta 2 |
| Itgb2l | integrin beta 2-like |
| Itgb3 | integrin beta 3 |
| Itgb8 | integrin beta 8 |
| Jup | junction plakoglobin |
| Lgals3 | lectin, galactose binding, soluble 3 |
| Lrg1 | leucine-rich alpha-2-glycoprotein 1 |
| Mir23a | microRNA 23a |
| Mir23b | microRNA 23b |
| Mir24-1 | microRNA 24-1 |
| Mir24-2 | microRNA 24-2 |
| Mir27a | microRNA 27a |
| Mir27b | microRNA 27b |
| Mmp9 | matrix metallopeptidase 9 |
| Mtdh | metadherin |
| Nfe2l2 | nuclear factor, erythroid derived 2, like 2 |
| Nodal | nodal |
| Nras | neuroblastoma ras oncogene |
| Ntrk1 | neurotrophic tyrosine kinase, receptor, type 1 |
| Pak4 | p21 (RAC1) activated kinase 4 |
| Pik3cd | phosphatidylinositol-4,5-bisphosphate 3-kinase catalytic subunit delta |
| Plcg1 | phospholipase C, gamma 1 |
| Prkcb | protein kinase C, beta |
| Prl2c2 | prolactin family 2, subfamily c, member 2 |
| Ptgis | prostaglandin I2 (prostacyclin) synthase |
| Pxn | paxillin |
| Rras | related RAS viral (r-ras) oncogene |
| Runx1 | runt related transcription factor 1 |
| Sash1 | SAM and SH3 domain containing 1 |
| Sfrp2 | secreted frizzled-related protein 2 |
| Smoc2 | SPARC related modular calcium binding 2 |
| Sp1 | trans-acting transcription factor 1 |
| Sphk1 | sphingosine kinase 1 |
| Stat3 | signal transducer and activator of transcription 3 |

| | |
|---|---|
| Stim1 | stromal interaction molecule 1 |
| Tbxa2r | thromboxane A2 receptor |
| Tert | telomerase reverse transcriptase |
| Tgfbr2 | transforming growth factor, beta receptor II |
| Tlr3 | toll-like receptor 3 |
| Tnfrsf1a | tumor necrosis factor receptor superfamily, member 1a |
| Uts2 | urotensin 2 |
| Uts2r | urotensin 2 receptor |
| Wars2 | tryptophanyl tRNA synthetase 2 (mitochondrial) |
| Wnk1 | WNK lysine deficient protein kinase 1 |
| Wnt5a | wingless-type MMTV integration site family, member 5A |
| Abl1 | c-abl oncogene 1, non-receptor tyrosine kinase |
| Mdk | midkine |
| Sirt6 | sirtuin 6 |
| Agtr1a | angiotensin II receptor, type 1a |
| Agtr1b | angiotensin II receptor, type 1b |
| Fgfbp1 | fibroblast growth factor binding protein 1 |
| Jcad | junctional cadherin 5 associated |
| Ppp1r16b | protein phosphatase 1, regulatory subunit 16B |
| Anxa1 | annexin A1 |
| Foxc2 | forkhead box C2 |
| Hdac7 | histone deacetylase 7 |
| Hdac9 | histone deacetylase 9 |
| Map3k3 | mitogen-activated protein kinase kinase kinase 3 |
| Pik3c2a | phosphatidylinositol-4-phosphate 3-kinase catalytic subunit type 2 alpha |
| Tgfbr3 | transforming growth factor, beta receptor III |
| Dll1 | delta like canonical Notch ligand 1 |
| Fut1 | fucosyltransferase 1 |
| Ghrl | ghrelin |
| Ghsr | growth hormone secretagogue receptor |
| Hmgb1 | high mobility group box 1 |
| Il10 | interleukin 10 |
| Jak1 | Janus kinase 1 |
| Jmjd8 | jumonji domain containing 8 |
| Pkm | pyruvate kinase, muscle |
| S100a1 | S100 calcium binding protein A1 |
| Slc39a12 | solute carrier family 39 (zinc transporter), member 12 |
| Smad1 | SMAD family member 1 |
| Tjp1 | tight junction protein 1 |
| Tnn | tenascin N |
| Cxcr4 | C-X-C motif chemokine receptor 4 |
| Bmper | BMP-binding endothelial regulator |
| Ccm2 | cerebral cavernous malformation 2 |

| | |
|---|---|
| Egln1 | egl-9 family hypoxia-inducible factor 1 |
| Emp2 | epithelial membrane protein 2 |
| Ets1 | E26 avian leukemia oncogene 1, 5' domain |
| Id1 | inhibitor of DNA binding 1, HLH protein |
| Mapk7 | mitogen-activated protein kinase 7 |
| Ppp1r15a | protein phosphatase 1, regulatory subunit 15A |
| Reck | reversion-inducing-cysteine-rich protein with kazal motifs |
| Rnh1 | ribonuclease/angiogenin inhibitor 1 |
| Sp100 | nuclear antigen Sp100 |
| Vash2 | vasohibin 2 |
| Fkbpl | FK506 binding protein-like |
| Fbxw7 | F-box and WD-40 domain protein 7 |
| Ceacam1 | CEA cell adhesion molecule 1 |
| Tspan18 | tetraspanin 18 |
| Cdc42 | cell division cycle 42 |
| Cdh13 | cadherin 13 |
| Clec14a | C-type lectin domain family 14, member a |
| E2f7 | E2F transcription factor 7 |
| E2f8 | E2F transcription factor 8 |
| Jmjd6 | jumonji domain containing 6 |
| Lef1 | lymphoid enhancer binding factor 1 |
| Loxl2 | lysyl oxidase-like 2 |
| Tgfb1 | transforming growth factor, beta 1 |
| Vstm4 | V-set and transmembrane domain containing 4 |
| Adipor2 | adiponectin receptor 2 |
| Npr2 | natriuretic peptide receptor 2 |

**Table S2. Clinical, laboratory, and demographic parameters of individuals with ADPKD and age/sex-matched controls – Pilot study.**

| **Subject Characteristics** | **Controls** | **ADPKD** | ***p-value*** |
|---|---|---|---|
| Number of subjects | 10 | 10 | N/A |
| Gender (Female/Male/NB) | 6/4/0 | 6/4/0 | N/A |
| Age (years) | 23.1 ± 3.1 | 22.7 ± 3.1 | N/A |
| Height (m) | 1.7 ± 0.1 | 1.7 ± 0.1 | 0.4897 |
| Weight (Kg) | 78.2 ± 13.0 | 84.9 ± 24.3 | 0.4458 |
| BSA ($m^2$) | 1.9 ± 0.2 | 2.0 ± 0.3 | 0.4575 |
| Systolic blood pressure (mmHg) | 114.6 ± 9.0 | 120.0 ± 7.4 | 0.1010 |
| Diastolic blood pressure (mmHg) | 70.2 ± 8.8 | 74.4 ± 6.9 | 0.2172 |
| Mean arterial pressure (mmHg) | 85.0 ± 7.9 | 89.6 ± 6.8 | 0.1375 |
| Serum creatinine (mg/dL) | 0.9 ± 0.1 | 0.8 ± 0.2 | 0.7052 |
| eGFR (ml/min/1.73$m^2$) | 109.3 ± 16.2 | 112.6 ± 11.7 | 0.5886 |
| Urine creatinine (mg/dL) | 179.4 ± 104.5 | 135.4 ± 81.9 | 0.2995 |
| Urine protein (mg/g creatinine) | 50.6 ± 43.1 | 48.0 ± 24.4 | 0.8829 |
| Urine pH | 6.3 ± 0.7 | 6.3 ± 0.5 | 0.9526 |
| Total Cholesterol (mg/dL) | 185.7 ± 24.7 | 186.1 ± 29.8 | 0.9774 |
| LDL cholesterol (mg/dL) | 117.6 ± 20.3 | 107.9 ± 35.8 | 0.5938 |
| HDL cholesterol (mg/dL) | 50.2 ± 7.7 | 50.5 ± 14.5 | 0.9674 |
| Triglycerides (mg/dL) | 122.6 ± 54.1 | 138.9 ± 79.9 | 0.6979 |
| Uric acid (mg/dL) | 5.6 ± 1.1 | 5.4 ± 1.7 | 0.8673 |
| htTKV (mL/m) | 175.5 ± 17.9 | 365.7 ± 108.3 | **0.0003** |
| Class A | N/A | 0 | N/A |
| Class B | N/A | 3 | N/A |
| Class C | N/A | 4 | N/A |
| Class D | N/A | 3 | N/A |
| Class E | N/A | 0 | N/A |

NB, non-binary; eGFR: estimated by CKD-EPI (2021); TKV: total kidney volume; htTKV: height-adjusted TKV;

| Table S3. Plasma metabolites | | |
|---|---|---|
| **Metabolite** | **Fold Difference** | **p.value** |
| 3-Methylhistidine | 0.96 | 0.3616 |
| Alanine | 1.19 | 0.1363 |
| allo-Isoleucine | 1.03 | 0.8205 |
| alpha-Aminoadipic-acid | 1.09 | 0.6466 |
| alpha-Amino-N-butyric-acid | 0.98 | 0.8333 |
| Arachidonic | 0.80 | 0.2776 |
| Arginine | 0.93 | 0.4291 |
| Asparagine | 0.93 | 0.2718 |
| Aspartic Acid | 0.94 | 0.6994 |
| beta-Alanine | 0.90 | 0.3672 |
| beta-Aminoisobutyric-acid | 0.86 | 0.6520 |
| Citrate | 0.92 | 0.3299 |
| Citrulline | 0.91 | 0.5921 |
| Cystathionine | 1.22 | 0.3880 |
| Cystine | 1.09 | 0.5030 |
| Docosahexaenoic Acid | 0.71 | 0.1011 |
| Elaidic | 0.78 | 0.3417 |
| Eicosapentaenoic Acid | 0.68 | 0.2174 |
| Ethanolamine | 0.95 | 0.5385 |
| Fumarate | 1.12 | 0.0563 |
| **gamma-Amino-N-butyric-acid** | **1.39** | **0.0296** |
| Glutamic Acid | 1.64 | 0.1187 |
| Glutamine | 1.05 | 0.2839 |
| Glycine | 1.12 | 0.3890 |
| Histidine | 0.95 | 0.5073 |
| **Homocysteine** | **3.27** | **0.0035** |
| Hydroxylysine | 1.22 | 0.3661 |
| Hydroxyproline | 1.18 | 0.3014 |
| Isoleucine | 1.07 | 0.5033 |
| alpha-Ketoglutarate | 0.97 | 0.4849 |
| Lactate | 1.08 | 0.1354 |
| Leucine | 1.04 | 0.6048 |
| Linoleic | 0.90 | 0.5990 |
| Linolenic | 0.90 | 0.5744 |
| Lysine | 1.03 | 0.7228 |
| Malate | 0.96 | 0.6002 |
| Methionine | 1.02 | 0.7260 |
| Myristic | 0.83 | 0.5175 |
| Oleic | 0.93 | 0.8138 |
| Ornithine | 1.17 | 0.1565 |
| Palmitic | 0.86 | 0.5758 |
| Palmitoleic | 0.88 | 0.6518 |

| | | |
|---|---|---|
| Phenylalanine | 1.01 | 0.9137 |
| Phosphoethanolamine | 0.99 | 0.9529 |
| Proline | 1.12 | 0.2353 |
| Sarcosine | 1.00 | 0.9870 |
| Serine | 1.03 | 0.7773 |
| Stearic | 0.89 | 0.4685 |
| Succinate | 1.05 | 0.1547 |
| Taurine | 0.90 | 0.3886 |
| Threonine | 0.93 | 0.5965 |
| Tryptophan | 0.97 | 0.6581 |
| Tyrosine | 0.97 | 0.7900 |
| Valine | 1.00 | 0.9709 |

13. Ebrahimi B, Eirin A, Li Z, et al. Mesenchymal stem cells improve medullary inflammation and fibrosis after revascularization of swine atherosclerotic renal artery stenosis. Research Support, N.I.H., Extramural

Research Support, Non-U.S. Gov't. *PLoS One*. 2013;8(7):e67474. doi:10.1371/journal.pone.0067474

45. JE J. *A user's guide to principal components*. Wiley Series in Probability and Statistics. John Wiley & Sons, Inc.; 1991.

### Figure S1

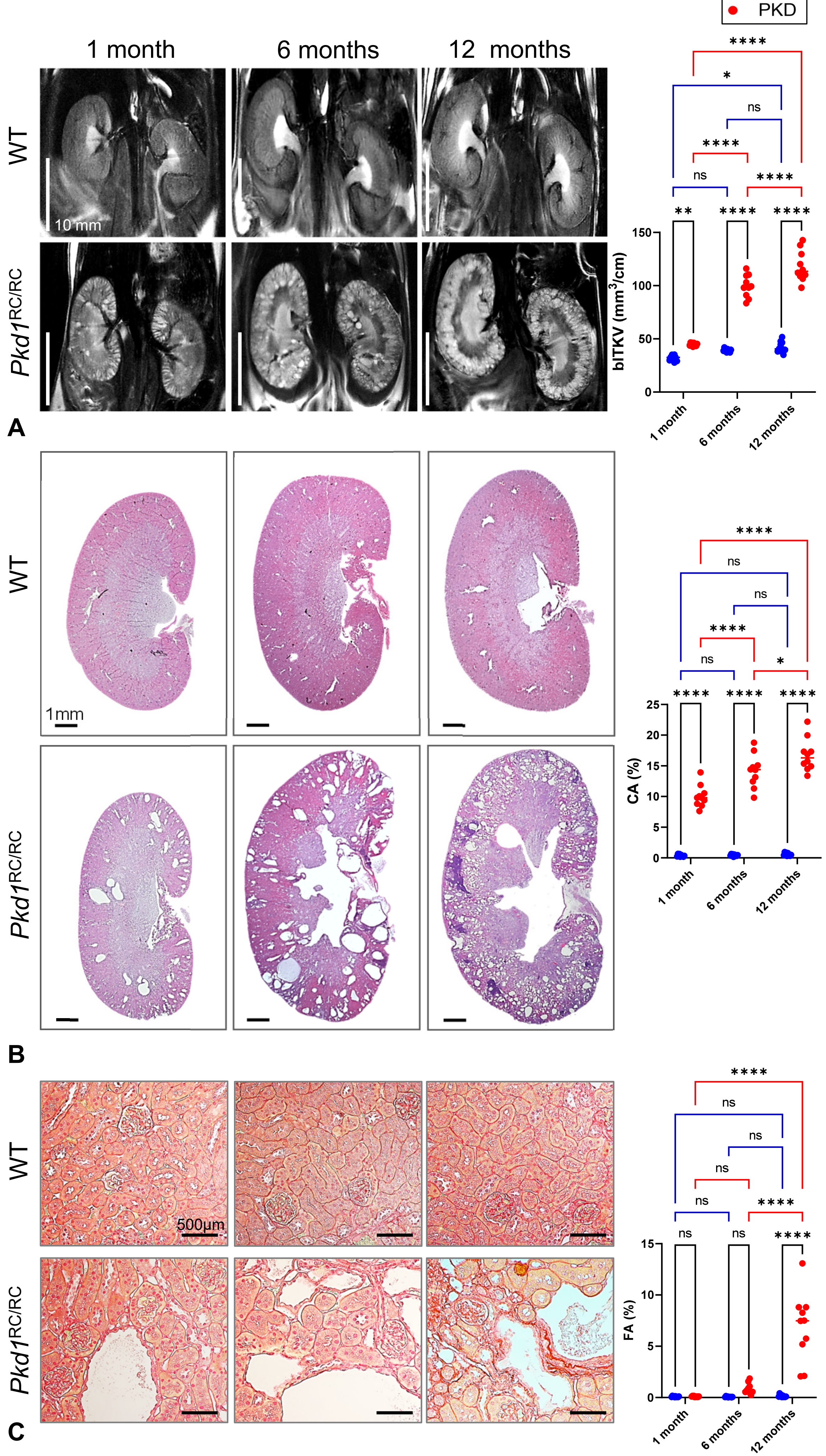

WT
PKD
1 month
6 months
12 months
WT
Pkd1$^{RC/RC}$
10 mm
A
blTKV (mm$^3$/cm)
150
100
50
0
**
****
ns
****
****
****
ns
*
****
1 month
6 months
12 months
WT
Pkd1$^{RC/RC}$
1mm
B
CA (%)
25
20
15
10
5
0
****
****
****
ns
*
****
ns
ns
ns
****
1 month
6 months
12 months
WT
Pkd1$^{RC/RC}$
500μm
C
FA (%)
15
10
5
0
ns
ns
****
ns
****
ns
ns
ns
****
1 month
6 months
12 months

Figure S2

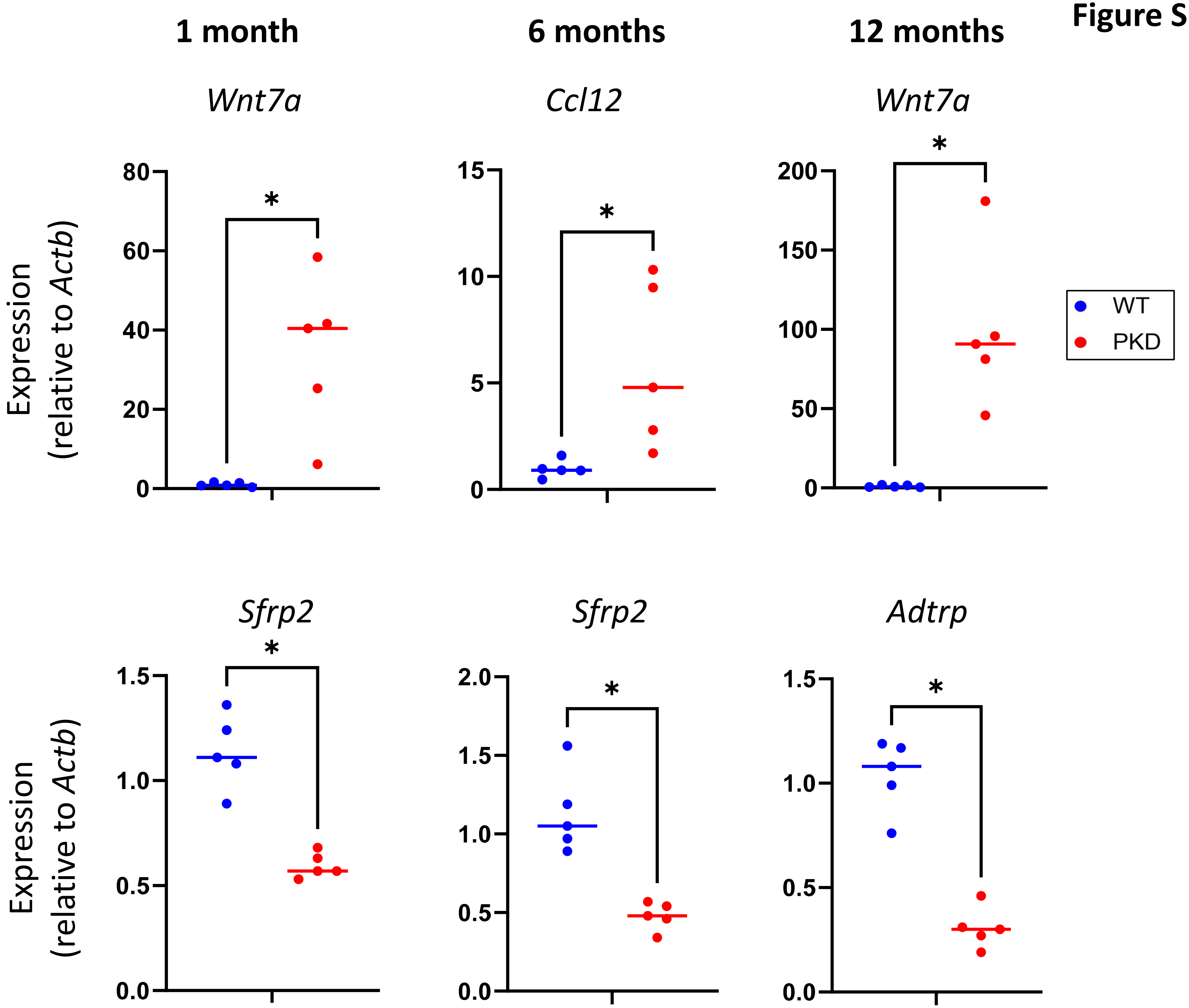

1 month
6 months
12 months
Wnt7a
Ccl12
Wnt7a
Sfrp2
Sfrp2
Adtrp
Expression (relative to Actb)
Expression (relative to Actb)
WT
PKD
*
80
60
40
20
0
15
10
5
0
200
150
100
50
0
1.5
1.0
0.5
0.0
2.0
1.5
1.0
0.5
0.0
1.5
1.0
0.5
0.0

**Figure S3**

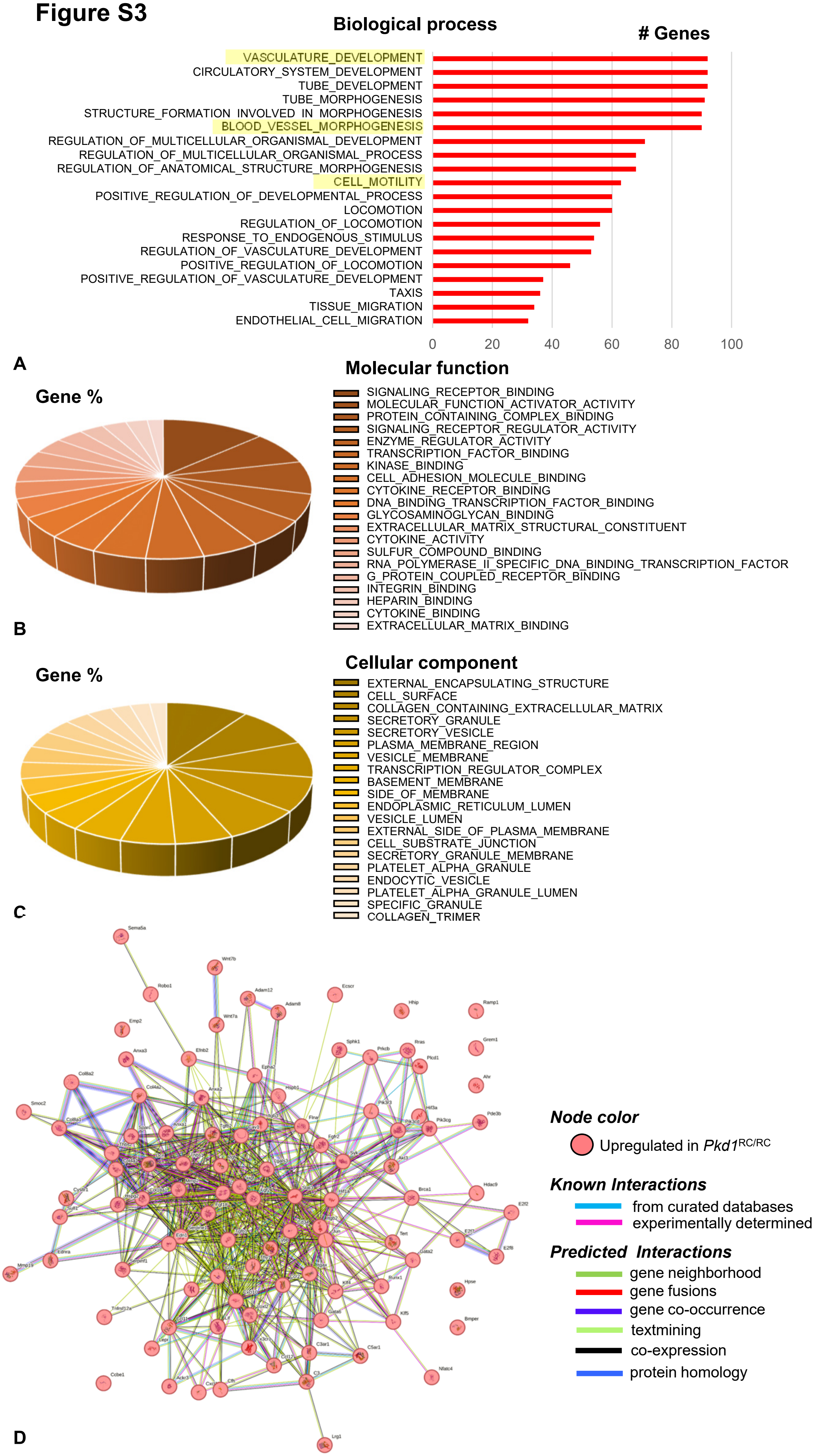

Biological process
### Genes
VASCULATURE_DEVELOPMENT
CIRCULATORY_SYSTEM_DEVELOPMENT
TUBE_DEVELOPMENT
TUBE_MORPHOGENESIS
STRUCTURE_FORMATION_INVOLVED_IN_MORPHOGENESIS
BLOOD_VESSEL_MORPHOGENESIS
REGULATION_OF_MULTICELLULAR_ORGANISMAL_DEVELOPMENT
REGULATION_OF_MULTICELLULAR_ORGANISMAL_PROCESS
REGULATION_OF_ANATOMICAL_STRUCTURE_MORPHOGENESIS
CELL_MOTILITY
POSITIVE_REGULATION_OF_DEVELOPMENTAL_PROCESS
LOCOMOTION
REGULATION_OF_LOCOMOTION
RESPONSE_TO_ENDOGENOUS_STIMULUS
REGULATION_OF_VASCULATURE_DEVELOPMENT
POSITIVE_REGULATION_OF_LOCOMOTION
POSITIVE_REGULATION_OF_VASCULATURE_DEVELOPMENT
TAXIS
TISSUE_MIGRATION
ENDOTHELIAL_CELL_MIGRATION
0
20
40
60
80
100
A
Molecular function
Gene %
SIGNALING_RECEPTOR_BINDING
MOLECULAR_FUNCTION_ACTIVATOR_ACTIVITY
PROTEIN_CONTAINING_COMPLEX_BINDING
SIGNALING_RECEPTOR_REGULATOR_ACTIVITY
ENZYME_REGULATOR_ACTIVITY
TRANSCRIPTION_FACTOR_BINDING
KINASE_BINDING
CELL_ADHESION_MOLECULE_BINDING
CYTOKINE_RECEPTOR_BINDING
DNA_BINDING_TRANSCRIPTION_FACTOR_BINDING
GLYCOSAMINOGLYCAN_BINDING
EXTRACELLULAR_MATRIX_STRUCTURAL_CONSTITUENT
CYTOKINE_ACTIVITY
SULFUR_COMPOUND_BINDING
RNA_POLYMERASE_II_SPECIFIC_DNA_BINDING_TRANSCRIPTION_FACTOR
G_PROTEIN_COUPLED_RECEPTOR_BINDING
INTEGRIN_BINDING
HEPARIN_BINDING
CYTOKINE_BINDING
EXTRACELLULAR_MATRIX_BINDING
B
Cellular component
Gene %
EXTERNAL_ENCAPSULATING_STRUCTURE
CELL_SURFACE
COLLAGEN_CONTAINING_EXTRACELLULAR_MATRIX
SECRETORY_GRANULE
SECRETORY_VESICLE
PLASMA_MEMBRANE_REGION
VESICLE_MEMBRANE
TRANSCRIPTION_REGULATOR_COMPLEX
BASEMENT_MEMBRANE
SIDE_OF_MEMBRANE
ENDOPLASMIC_RETICULUM_LUMEN
VESICLE_LUMEN
EXTERNAL_SIDE_OF_PLASMA_MEMBRANE
CELL_SUBSTRATE_JUNCTION
SECRETORY_GRANULE_MEMBRANE
PLATELET_ALPHA_GRANULE
ENDOCYTIC_VESICLE
PLATELET_ALPHA_GRANULE_LUMEN
SPECIFIC_GRANULE
COLLAGEN_TRIMER
C
Node color
Upregulated in Pkd1RC/RC
Known Interactions
from curated databases
experimentally determined
Predicted Interactions
gene neighborhood
gene fusions
gene co-occurrence
textmining
co-expression
protein homology
D

Figure S4

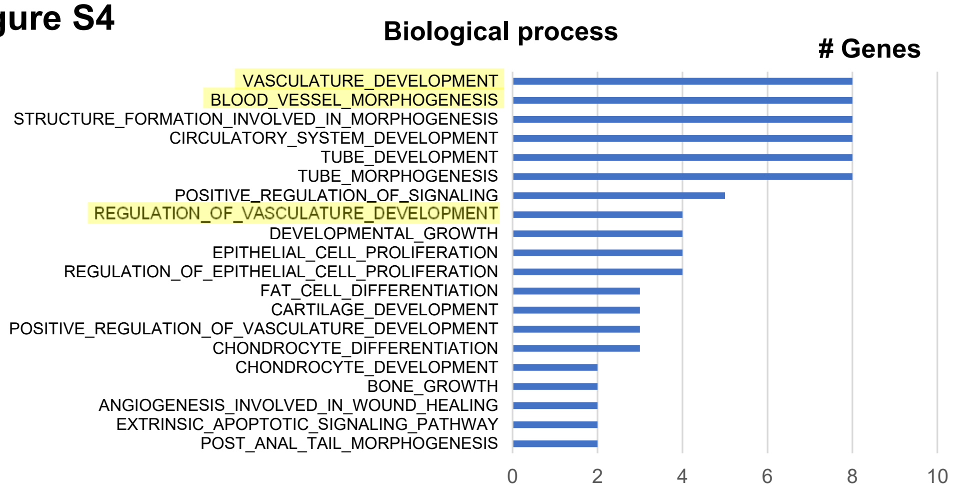


A

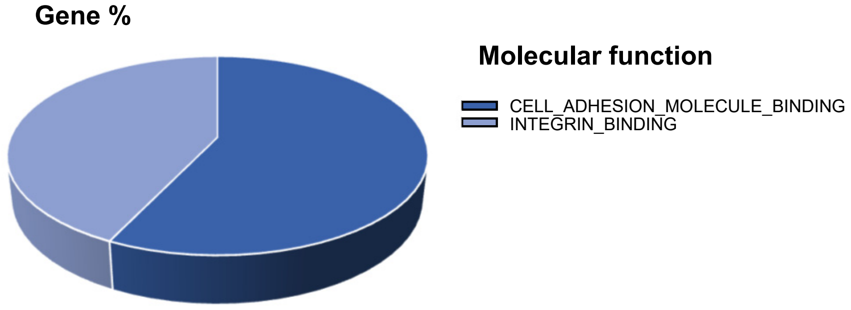


B

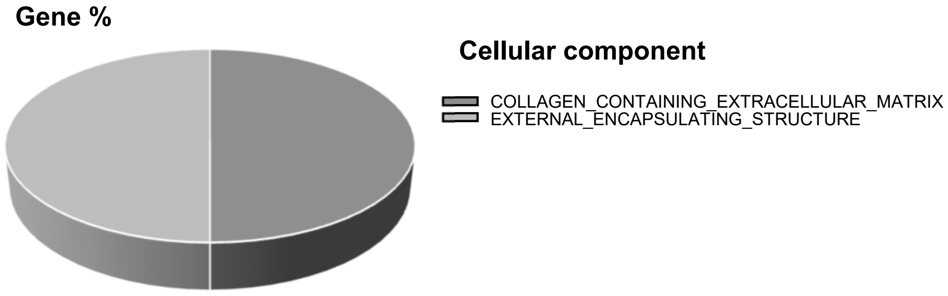


C

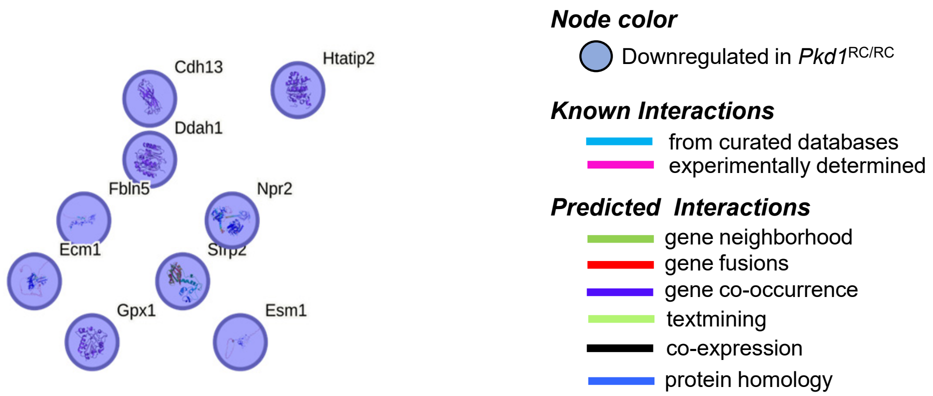


D

#### Figure S5

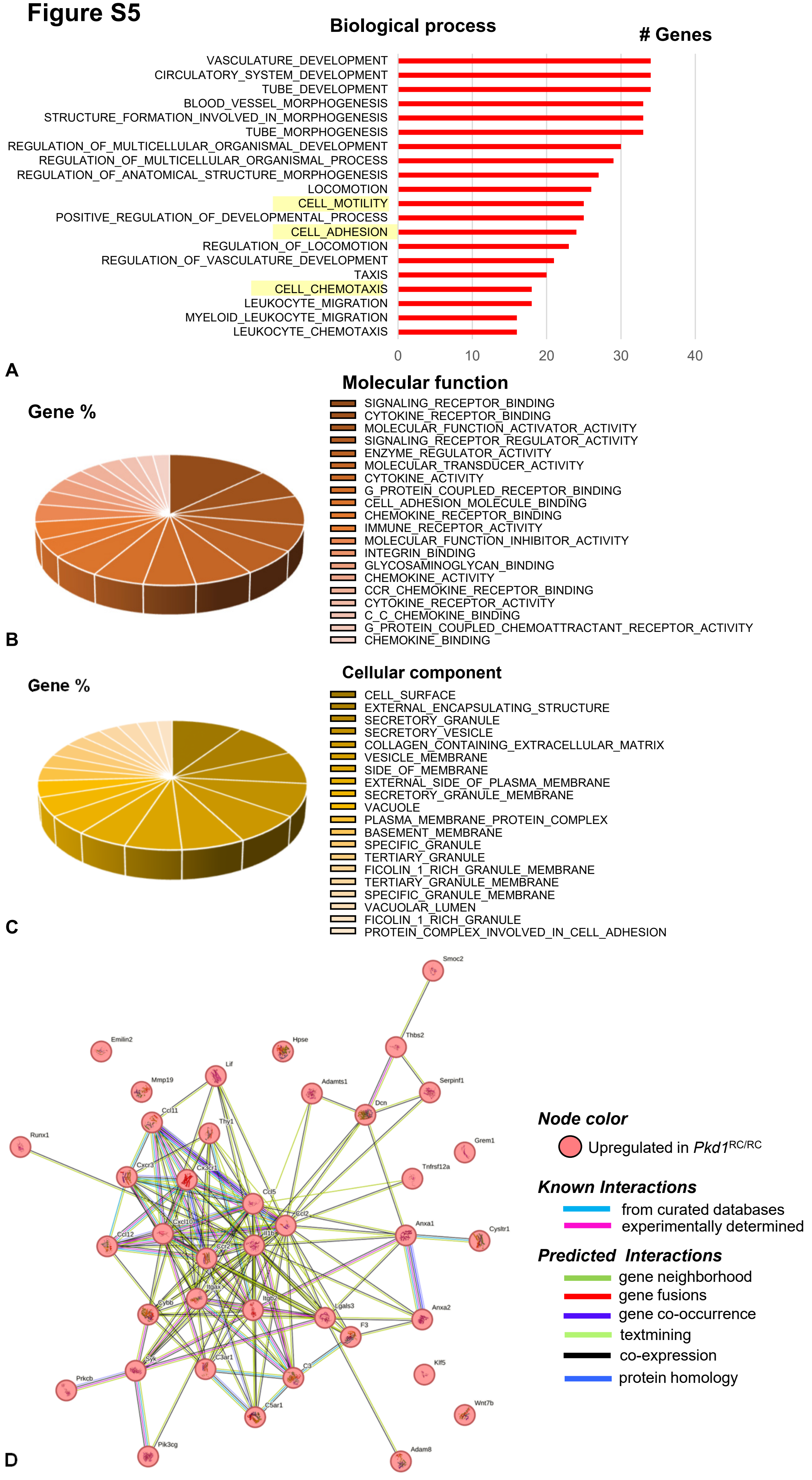
Biological process
### Genes
VASCULATURE_DEVELOPMENT
CIRCULATORY_SYSTEM_DEVELOPMENT
TUBE_DEVELOPMENT
BLOOD_VESSEL_MORPHOGENESIS
STRUCTURE_FORMATION_INVOLVED_IN_MORPHOGENESIS
TUBE_MORPHOGENESIS
REGULATION_OF_MULTICELLULAR_ORGANISMAL_DEVELOPMENT
REGULATION_OF_MULTICELLULAR_ORGANISMAL_PROCESS
REGULATION_OF_ANATOMICAL_STRUCTURE_MORPHOGENESIS
LOCOMOTION
CELL_MOTILITY
POSITIVE_REGULATION_OF_DEVELOPMENTAL_PROCESS
CELL_ADHESION
REGULATION_OF_LOCOMOTION
REGULATION_OF_VASCULATURE_DEVELOPMENT
TAXIS
CELL_CHEMOTAXIS
LEUKOCYTE_MIGRATION
MYELOID_LEUKOCYTE_MIGRATION
LEUKOCYTE_CHEMOTAXIS
0
10
20
30
40
A
Molecular function
Gene %
SIGNALING_RECEPTOR_BINDING
CYTOKINE_RECEPTOR_BINDING
MOLECULAR_FUNCTION_ACTIVATOR_ACTIVITY
SIGNALING_RECEPTOR_REGULATOR_ACTIVITY
ENZYME_REGULATOR_ACTIVITY
MOLECULAR_TRANSDUCER_ACTIVITY
CYTOKINE_ACTIVITY
G_PROTEIN_COUPLED_RECEPTOR_BINDING
CELL_ADHESION_MOLECULE_BINDING
CHEMOKINE_RECEPTOR_BINDING
IMMUNE_RECEPTOR_ACTIVITY
MOLECULAR_FUNCTION_INHIBITOR_ACTIVITY
INTEGRIN_BINDING
GLYCOSAMINOGLYCAN_BINDING
CHEMOKINE_ACTIVITY
CCR_CHEMOKINE_RECEPTOR_BINDING
CYTOKINE_RECEPTOR_ACTIVITY
C_C_CHEMOKINE_BINDING
G_PROTEIN_COUPLED_CHEMOATTRACTANT_RECEPTOR_ACTIVITY
CHEMOKINE_BINDING
B
Cellular component
Gene %
CELL_SURFACE
EXTERNAL_ENCAPSULATING_STRUCTURE
SECRETORY_GRANULE
SECRETORY_VESICLE
COLLAGEN_CONTAINING_EXTRACELLULAR_MATRIX
VESICLE_MEMBRANE
SIDE_OF_MEMBRANE
EXTERNAL_SIDE_OF_PLASMA_MEMBRANE
SECRETORY_GRANULE_MEMBRANE
VACUOLE
PLASMA_MEMBRANE_PROTEIN_COMPLEX
BASEMENT_MEMBRANE
SPECIFIC_GRANULE
TERTIARY_GRANULE
FICOLIN_1_RICH_GRANULE_MEMBRANE
TERTIARY_GRANULE_MEMBRANE
SPECIFIC_GRANULE_MEMBRANE
VACUOLAR_LUMEN
FICOLIN_1_RICH_GRANULE
PROTEIN_COMPLEX_INVOLVED_IN_CELL_ADHESION
C
Smoc2
Emilin2
Hpse
Thbs2
Lif
Mmp19
Adamts1
Serpinf1
Dcn
Ccl11
Thy1
Runx1
Grem1
Tnfrsf12a
Cxcr3
Cx3cr1
Ccl5
Ccl2
Anxa1
Cysltr1
Ccl12
Cxcl10
Il1b
Ccr2
Itgax
Cybb
Itgb2
Lgals3
Anxa2
F3
C3ar1
C3
Syk
Klf5
Prkcb
C5ar1
Wnt7b
Pik3cg
Adam8
Node color
Upregulated in Pkd1RC/RC
Known Interactions
from curated databases
experimentally determined
Predicted Interactions
gene neighborhood
gene fusions
gene co-occurrence
textmining
co-expression
protein homology
D

#### Figure S6

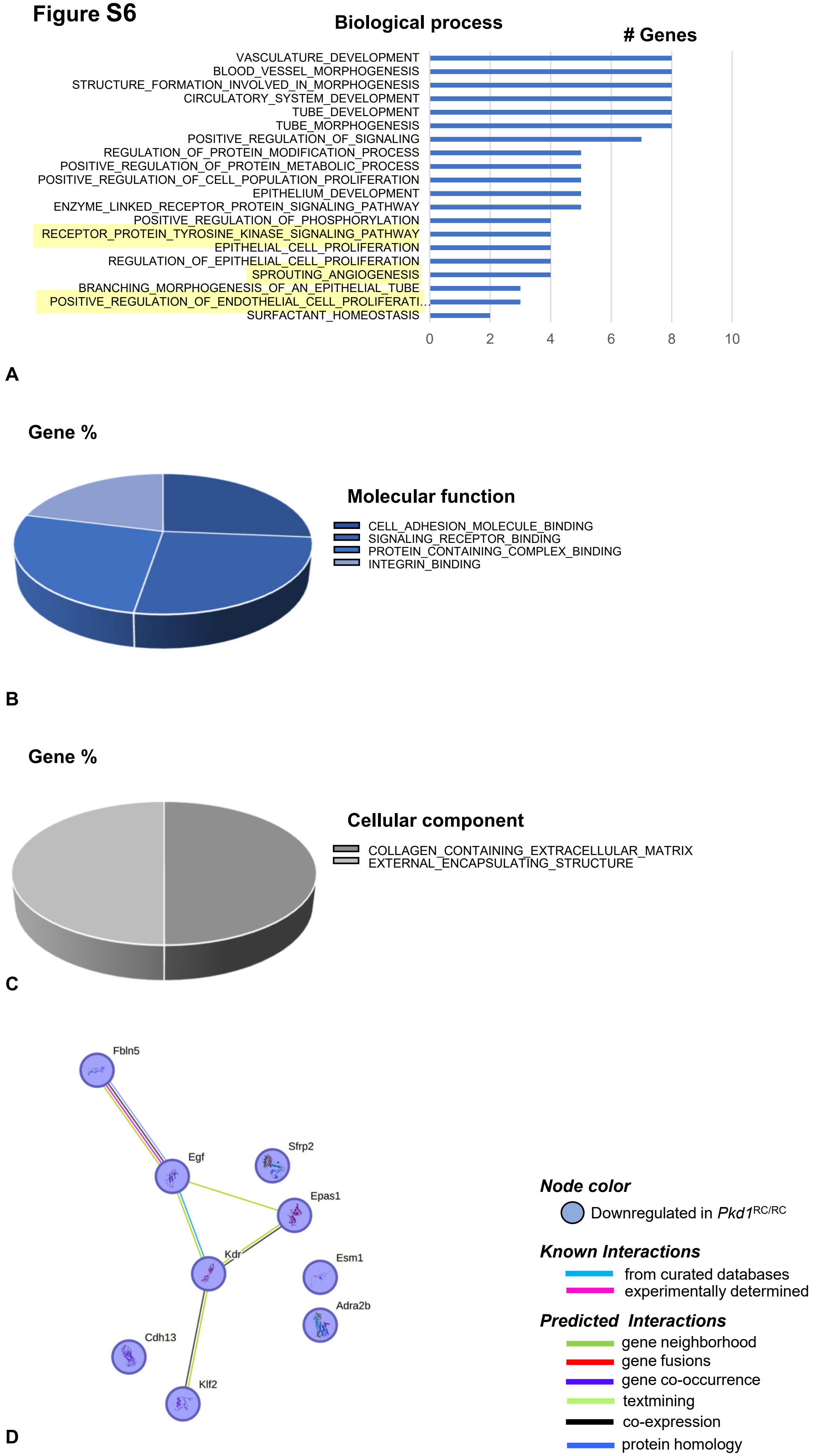
Biological process
### Genes
VASCULATURE_DEVELOPMENT
BLOOD_VESSEL_MORPHOGENESIS
STRUCTURE_FORMATION_INVOLVED_IN_MORPHOGENESIS
CIRCULATORY_SYSTEM_DEVELOPMENT
TUBE_DEVELOPMENT
TUBE_MORPHOGENESIS
POSITIVE_REGULATION_OF_SIGNALING
REGULATION_OF_PROTEIN_MODIFICATION_PROCESS
POSITIVE_REGULATION_OF_PROTEIN_METABOLIC_PROCESS
POSITIVE_REGULATION_OF_CELL_POPULATION_PROLIFERATION
EPITHELIUM_DEVELOPMENT
ENZYME_LINKED_RECEPTOR_PROTEIN_SIGNALING_PATHWAY
POSITIVE_REGULATION_OF_PHOSPHORYLATION
RECEPTOR_PROTEIN_TYROSINE_KINASE_SIGNALING_PATHWAY
EPITHELIAL_CELL_PROLIFERATION
REGULATION_OF_EPITHELIAL_CELL_PROLIFERATION
SPROUTING_ANGIOGENESIS
BRANCHING_MORPHOGENESIS_OF_AN_EPITHELIAL_TUBE
POSITIVE_REGULATION_OF_ENDOTHELIAL_CELL_PROLIFERATI…
SURFACTANT_HOMEOSTASIS
0
2
4
6
8
10
A
Gene %
Molecular function
CELL_ADHESION_MOLECULE_BINDING
SIGNALING_RECEPTOR_BINDING
PROTEIN_CONTAINING_COMPLEX_BINDING
INTEGRIN_BINDING
B
Gene %
Cellular component
COLLAGEN_CONTAINING_EXTRACELLULAR_MATRIX
EXTERNAL_ENCAPSULATING_STRUCTURE
C
Fbln5
Egf
Sfrp2
Epas1
Kdr
Esm1
Adra2b
Cdh13
Klf2
Node color
Downregulated in Pkd1RC/RC
Known Interactions
from curated databases
experimentally determined
Predicted Interactions
gene neighborhood
gene fusions
gene co-occurrence
textmining
co-expression
protein homology
D

**Figure S7**

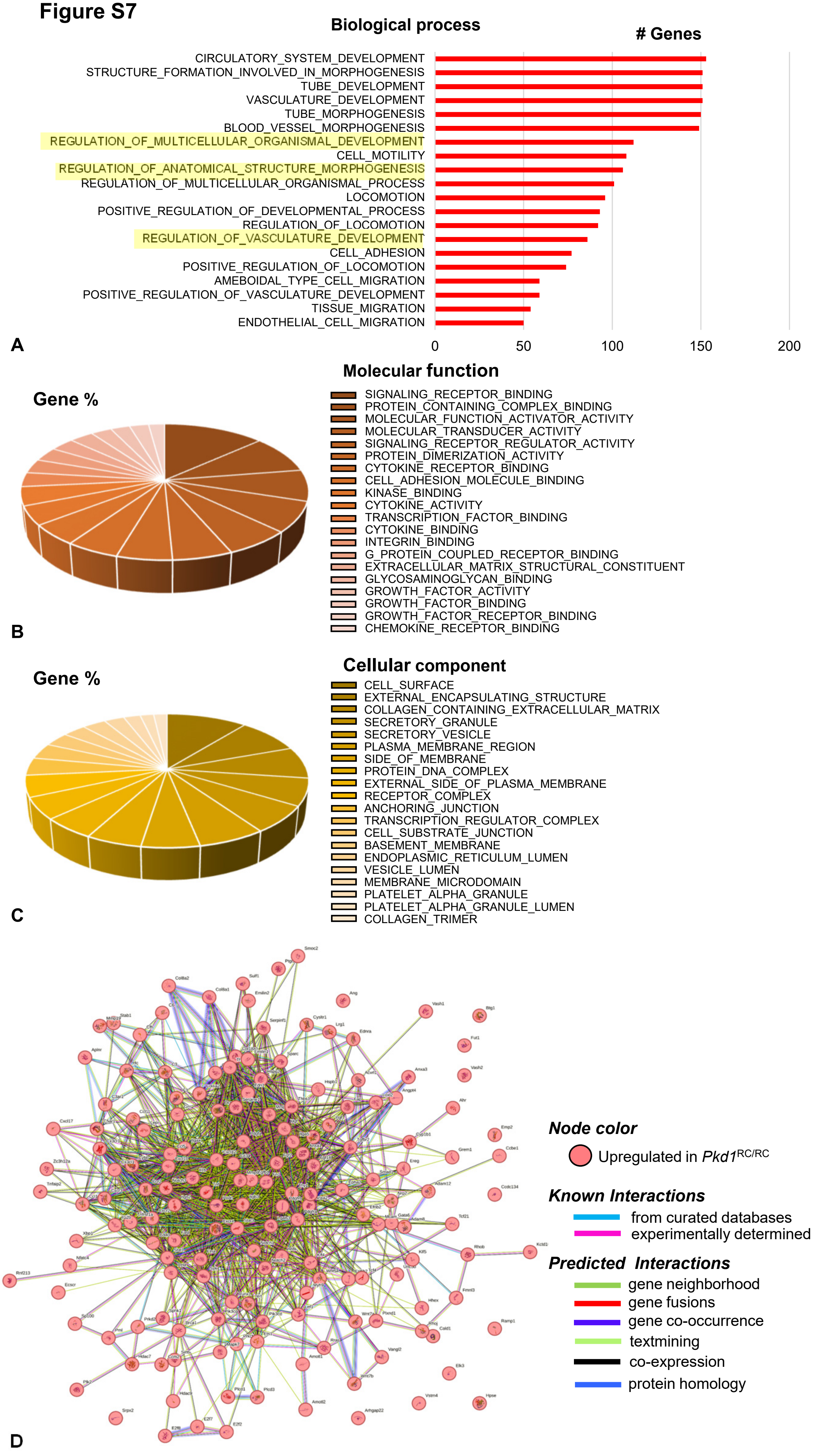

Biological process
### Genes
CIRCULATORY_SYSTEM_DEVELOPMENT
STRUCTURE_FORMATION_INVOLVED_IN_MORPHOGENESIS
TUBE_DEVELOPMENT
VASCULATURE_DEVELOPMENT
TUBE_MORPHOGENESIS
BLOOD_VESSEL_MORPHOGENESIS
REGULATION_OF_MULTICELLULAR_ORGANISMAL_DEVELOPMENT
CELL_MOTILITY
REGULATION_OF_ANATOMICAL_STRUCTURE_MORPHOGENESIS
REGULATION_OF_MULTICELLULAR_ORGANISMAL_PROCESS
LOCOMOTION
POSITIVE_REGULATION_OF_DEVELOPMENTAL_PROCESS
REGULATION_OF_LOCOMOTION
REGULATION_OF_VASCULATURE_DEVELOPMENT
CELL_ADHESION
POSITIVE_REGULATION_OF_LOCOMOTION
AMEBOIDAL_TYPE_CELL_MIGRATION
POSITIVE_REGULATION_OF_VASCULATURE_DEVELOPMENT
TISSUE_MIGRATION
ENDOTHELIAL_CELL_MIGRATION
0
50
100
150
200
A
Molecular function
Gene %
SIGNALING_RECEPTOR_BINDING
PROTEIN_CONTAINING_COMPLEX_BINDING
MOLECULAR_FUNCTION_ACTIVATOR_ACTIVITY
MOLECULAR_TRANSDUCER_ACTIVITY
SIGNALING_RECEPTOR_REGULATOR_ACTIVITY
PROTEIN_DIMERIZATION_ACTIVITY
CYTOKINE_RECEPTOR_BINDING
CELL_ADHESION_MOLECULE_BINDING
KINASE_BINDING
CYTOKINE_ACTIVITY
TRANSCRIPTION_FACTOR_BINDING
CYTOKINE_BINDING
INTEGRIN_BINDING
G_PROTEIN_COUPLED_RECEPTOR_BINDING
EXTRACELLULAR_MATRIX_STRUCTURAL_CONSTITUENT
GLYCOSAMINOGLYCAN_BINDING
GROWTH_FACTOR_ACTIVITY
GROWTH_FACTOR_BINDING
GROWTH_FACTOR_RECEPTOR_BINDING
CHEMOKINE_RECEPTOR_BINDING
B
Cellular component
Gene %
CELL_SURFACE
EXTERNAL_ENCAPSULATING_STRUCTURE
COLLAGEN_CONTAINING_EXTRACELLULAR_MATRIX
SECRETORY_GRANULE
SECRETORY_VESICLE
PLASMA_MEMBRANE_REGION
SIDE_OF_MEMBRANE
PROTEIN_DNA_COMPLEX
EXTERNAL_SIDE_OF_PLASMA_MEMBRANE
RECEPTOR_COMPLEX
ANCHORING_JUNCTION
TRANSCRIPTION_REGULATOR_COMPLEX
CELL_SUBSTRATE_JUNCTION
BASEMENT_MEMBRANE
ENDOPLASMIC_RETICULUM_LUMEN
VESICLE_LUMEN
MEMBRANE_MICRODOMAIN
PLATELET_ALPHA_GRANULE
PLATELET_ALPHA_GRANULE_LUMEN
COLLAGEN_TRIMER
C
Node color
Upregulated in Pkd1 RC/RC
Known Interactions
from curated databases
experimentally determined
Predicted Interactions
gene neighborhood
gene fusions
gene co-occurrence
textmining
co-expression
protein homology
D

**Figure S8**

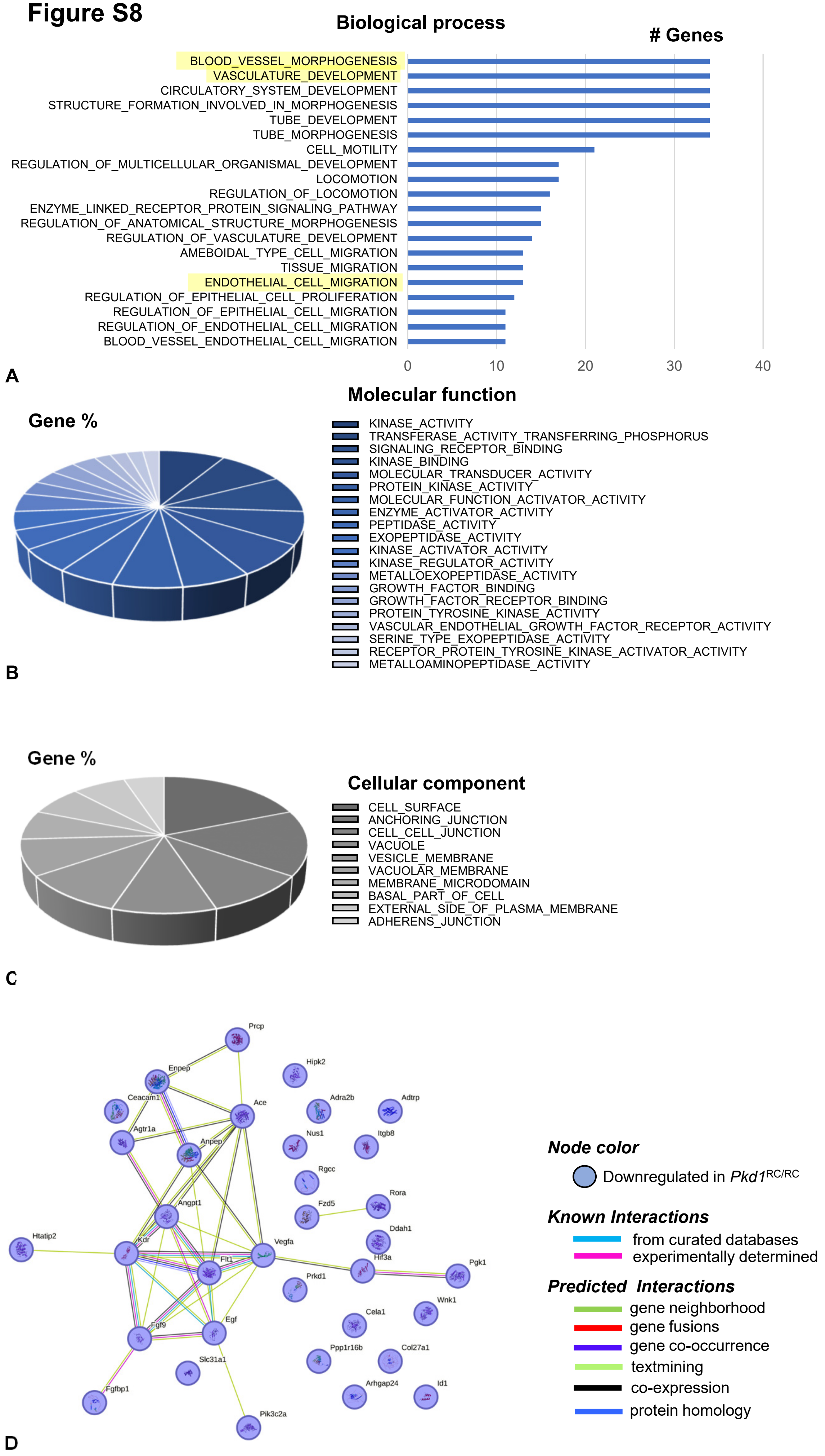

Biological process
### Genes
BLOOD_VESSEL_MORPHOGENESIS
VASCULATURE_DEVELOPMENT
CIRCULATORY_SYSTEM_DEVELOPMENT
STRUCTURE_FORMATION_INVOLVED_IN_MORPHOGENESIS
TUBE_DEVELOPMENT
TUBE_MORPHOGENESIS
CELL_MOTILITY
REGULATION_OF_MULTICELLULAR_ORGANISMAL_DEVELOPMENT
LOCOMOTION
REGULATION_OF_LOCOMOTION
ENZYME_LINKED_RECEPTOR_PROTEIN_SIGNALING_PATHWAY
REGULATION_OF_ANATOMICAL_STRUCTURE_MORPHOGENESIS
REGULATION_OF_VASCULATURE_DEVELOPMENT
AMEBOIDAL_TYPE_CELL_MIGRATION
TISSUE_MIGRATION
ENDOTHELIAL_CELL_MIGRATION
REGULATION_OF_EPITHELIAL_CELL_PROLIFERATION
REGULATION_OF_EPITHELIAL_CELL_MIGRATION
REGULATION_OF_ENDOTHELIAL_CELL_MIGRATION
BLOOD_VESSEL_ENDOTHELIAL_CELL_MIGRATION
0
10
20
30
40
A
Molecular function
Gene %
KINASE_ACTIVITY
TRANSFERASE_ACTIVITY_TRANSFERRING_PHOSPHORUS
SIGNALING_RECEPTOR_BINDING
KINASE_BINDING
MOLECULAR_TRANSDUCER_ACTIVITY
PROTEIN_KINASE_ACTIVITY
MOLECULAR_FUNCTION_ACTIVATOR_ACTIVITY
ENZYME_ACTIVATOR_ACTIVITY
PEPTIDASE_ACTIVITY
EXOPEPTIDASE_ACTIVITY
KINASE_ACTIVATOR_ACTIVITY
KINASE_REGULATOR_ACTIVITY
METALLOEXOPEPTIDASE_ACTIVITY
GROWTH_FACTOR_BINDING
GROWTH_FACTOR_RECEPTOR_BINDING
PROTEIN_TYROSINE_KINASE_ACTIVITY
VASCULAR_ENDOTHELIAL_GROWTH_FACTOR_RECEPTOR_ACTIVITY
SERINE_TYPE_EXOPEPTIDASE_ACTIVITY
RECEPTOR_PROTEIN_TYROSINE_KINASE_ACTIVATOR_ACTIVITY
METALLOAMINOPEPTIDASE_ACTIVITY
B
Gene %
Cellular component
CELL_SURFACE
ANCHORING_JUNCTION
CELL_CELL_JUNCTION
VACUOLE
VESICLE_MEMBRANE
VACUOLAR_MEMBRANE
MEMBRANE_MICRODOMAIN
BASAL_PART_OF_CELL
EXTERNAL_SIDE_OF_PLASMA_MEMBRANE
ADHERENS_JUNCTION
C
Prcp
Hipk2
Enpep
Ceacam1
Ace
Adra2b
Adtrp
Agtr1a
Anpep
Nus1
Itgb8
Rgcc
Angpt1
Fzd5
Rora
Ddah1
Htatip2
Kdr
Vegfa
Flt1
Hif3a
Pgk1
Prkd1
Cela1
Wnk1
Egf
Fgf9
Slc31a1
Ppp1r16b
Col27a1
Fgfbp1
Arhgap24
Id1
Pik3c2a
D
Node color
Downregulated in Pkd1RC/RC
Known Interactions
from curated databases
experimentally determined
Predicted Interactions
gene neighborhood
gene fusions
gene co-occurrence
textmining
co-expression
protein homology

Figure S9

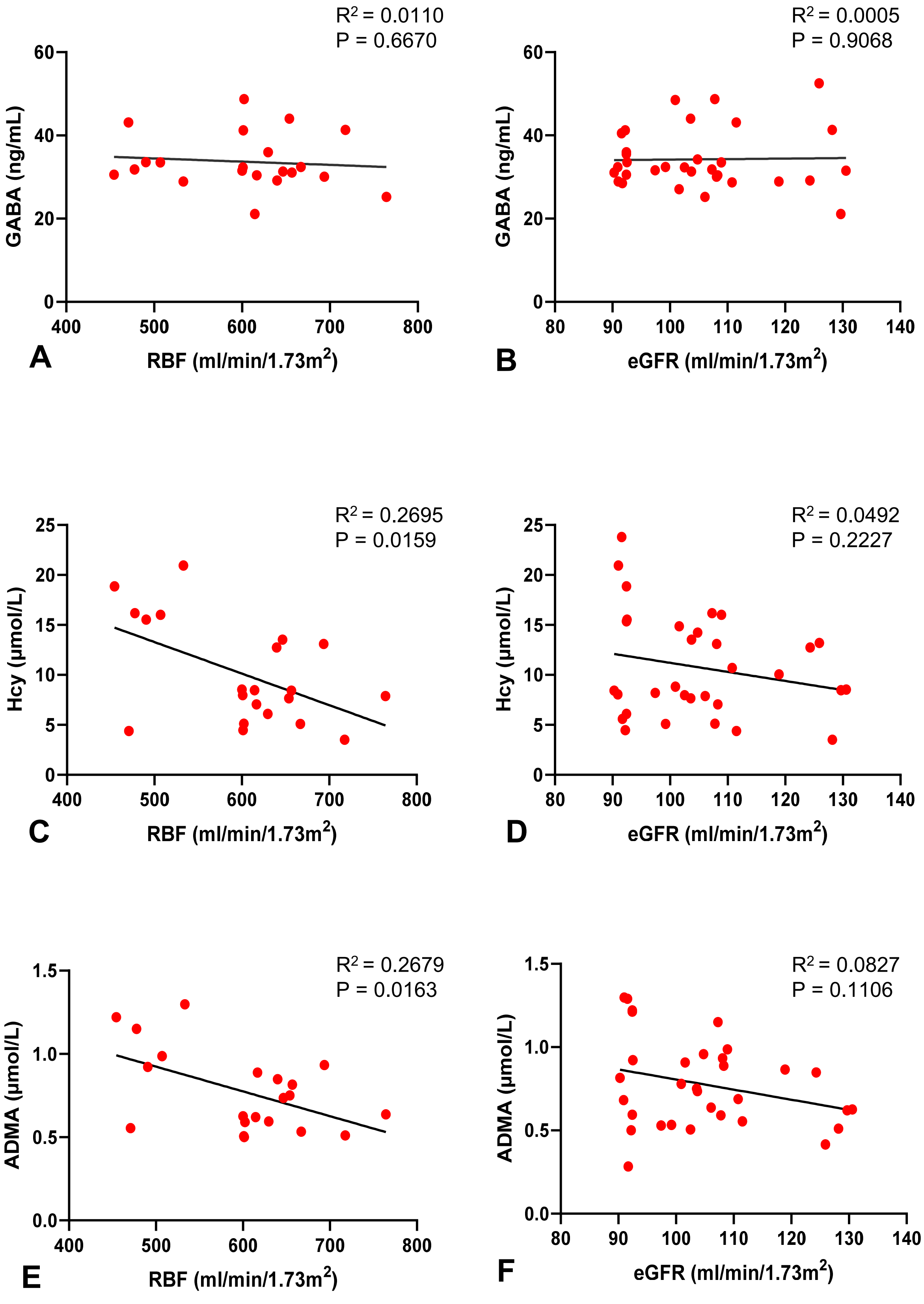

R2 = 0.0110
P = 0.6670
GABA (ng/mL)
RBF (ml/min/1.73m2)
A
R2 = 0.0005
P = 0.9068
GABA (ng/mL)
eGFR (ml/min/1.73m2)
B
R2 = 0.2695
P = 0.0159
Hcy (µmol/L)
RBF (ml/min/1.73m2)
C
R2 = 0.0492
P = 0.2227
Hcy (µmol/L)
eGFR (ml/min/1.73m2)
D
R2 = 0.2679
P = 0.0163
ADMA (µmol/L)
RBF (ml/min/1.73m2)
E
R2 = 0.0827
P = 0.1106
ADMA (µmol/L)
eGFR (ml/min/1.73m2)
F

#### SUPPLEMENTAL FIGURE LEGENDS

***Supplemental Figure 1.*** Kidney disease progression in *Pkd1*$^{RC/RC}$. Representative magnetic resonance imaging (MRI) of kidney coronal sections from live *Pkd1*$^{RC/RC}$ and WT mice at 1, 6, and 12 months and quantification of total kidney volume (TKV) adjusted by body length (bl) **(A)**. Representative kidney sections of 1-, 6-, and 12-month *Pkd1*$^{RC/RC}$ and WT mice stained with Hematoxylin and Eosin (H&E) **(B)** and picrosirius red **(C)**, and quantification of cystic (CA) and fibrotic areas (FA) (n=5M and 5F per genotype for each time point). *p<0.05, **p<0.01, ****p<0.0001 compared to WT at each time point.

***Supplemental Figure 2.*** Validation of mRNA-seq analysis. *Expression levels (real-time qPCR) of Wnt7a and Sfrp2, Ccl12 and Sfrp2, and Wnt7a* and *Adtrp in Pkd1*$^{RC/RC}$ *compared to WT kidneys* at 1, 6, and 12 months respectively (n=3M and 2F WT and n=2M and n=3F *Pkd1*$^{RC/RC}$ for each time point). *p<0.05 compared to WT at each time point. mRNA expression was normalized to β-actin (*Actb*).

***Supplemental Figure 3.*** *Vasculature-related genes upregulated in Pkd1*$^{RC/RC}$ *kidneys at 1 month*. Gene set enrichment analysis (GSEA) analysis of vasculature-related genes upregulated in *Pkd1*$^{RC/RC}$ compared to WT kidneys at 1 month and their classification by biological process **(A)**, molecular function **(B)**, and cellular component **(C)**. Clusters of physical and functional protein interactions among vasculature-related genes upregulated in *Pkd1*$^{RC/RC}$ at 1 month (higher strength of interaction represented by multiple strings between the genes shown in different colors) **(D)**.

***Supplemental Figure 4.*** *Vasculature-related genes downregulated in Pkd1$^{RC/RC}$ kidneys at 1 month*. Gene set enrichment analysis (GSEA) analysis of vasculature-related genes downregulated in *Pkd1*$^{RC/RC}$ compared to WT kidneys at 1 month and their classification by cellular component **(A)**, molecular function **(B)**, and biological process **(C)**. Clusters of physical and functional protein interactions among vasculature-related genes downregulated in *Pkd1*$^{RC/RC}$ at 1 month (higher strength of interaction represented by multiple strings between the genes shown in different colors) **(D)**.

***Supplemental Figure 5.*** *Vasculature-related genes upregulated in Pkd1$^{RC/RC}$ kidneys at 6 months*. Gene set enrichment analysis (GSEA) analysis of vasculature-related genes upregulated in *Pkd1*$^{RC/RC}$ compared to WT kidneys at 6 months and their classification by cellular component **(A)**, molecular function **(B)**, and biological process **(C)**. Clusters of physical and functional protein interactions among vasculature-related genes upregulated in *Pkd1*$^{RC/RC}$ at 6 months (higher strength of interaction represented by multiple strings between the genes shown in different colors) **(D)**.

***Supplemental Figure 6.*** *Vasculature-related genes downregulated in Pkd1$^{RC/RC}$ kidneys at 6 months*. Gene set enrichment analysis (GSEA) analysis of vasculature-related genes downregulated in *Pkd1*$^{RC/RC}$ compared to WT kidneys at 6 months and their classification by cellular component **(A)**, molecular function **(B)**, and biological process **(C)**. Clusters of physical and functional protein interactions among vasculature-related genes downregulated in *Pkd1*$^{RC/RC}$ at 6 months (higher strength of interaction represented by multiple strings between the genes shown in different colors) **(D)**.

***Supplemental Figure 7.*** *Vasculature-related genes upregulated in Pkd1$^{RC/RC}$ kidneys at 12 months*. Gene set enrichment analysis (GSEA) analysis of vasculature-reated genes upregulated in *Pkd1*$^{RC/RC}$ compared to WT kidneys at 12 months and their classification by cellular component **(A)**, molecular function **(B)**, and biological process **(C)**. Clusters of physical and functional protein interactions among vasculature-related genes upregulated in *Pkd1*$^{RC/RC}$ at 12 months (higher strength of interaction represented by multiple strings between the genes shown in different colors) **(D)**.

***Supplemental Figure 8.*** *Vasculature-related genes downregulated in Pkd1*$^{RC/RC}$ *kidneys at 12 months*. Gene set enrichment analysis (GSEA) analysis of vasculature-related genes downregulated in *Pkd1*$^{RC/RC}$ compared to WT kidneys at 12 months and their classification by cellular component **(A)**, molecular function **(B)**, and biological process **(C)**. Clusters of physical and functional protein interactions among vasculature-related genes downregulated in *Pkd1*$^{RC/RC}$ at 12 months (higher strength of interaction represented by multiple strings between the genes shown in different colors) **(D)**.

***Supplemental Figure 9.*** *Plasma gamma-aminobutyric acid (GABA), homocysteine (Hcy), and asymmetric dimethylarginine (ADMA) and their correlation RBF and eGFR.* Correlations between GABA and renal blood flow (RBF) **(A)**, GABA and estimated glomerular filtration rate (eGFR) **(B)**, Hcy and RBF **(C)**, Hcy and eGFR **(D)**, ADMA and RBF **(E)**, and ADMA and eGFR **(F)**.
